# Ectopic histone clipping in the mouse model of progressive myoclonus epilepsy

**DOI:** 10.1101/2020.08.21.261180

**Authors:** Eduard Daura, Saara Tegelberg, Masahito Yoshihara, Christopher Jackson, Francesca Simonetti, Katri Aksentjeff, Sini Ezer, Paula Hakala, Shintaro Katayama, Juha Kere, Anna-Elina Lehesjoki, Tarja Joensuu

## Abstract

We establish cystatin B (CSTB) as a regulator of histone H3 tail clipping in murine neural progenitor cells (NPCs) and provide evidence suggesting that epigenetic dysregulation contributes to the early pathogenesis in brain disorders associated with deficient CSTB function. We show that NPCs undergo regulated cleavage of the N-terminal tail of histone H3 at threonine 22 (H3T22) transiently upon induction of differentiation. CSTB-deficient NPCs present premature activation of H3T22 clipping during self-renewal mediated by increased activity of cathepsins L and B. During differentiation, the proportion of immature committed neurons undergoing H3T22 clipping is significantly higher in CSTB-deficient than in wild-type NPCs, with no observable decline within 12 days post-differentiation. CSTB-deficient NPCs exhibit significant transcriptional changes highlighting altered expression of nuclear-encoded mitochondrial genes. These changes are associated with significantly impaired respiratory capacity of differentiating NPCs devoid of CSTB. Our data expand the mechanistic understanding of diseases associated with CSTB deficiency.

## INTRODUCTION

Progressive myoclonus epilepsy, EPM1 (Unverricht-Lundborg disease; OMIM 254800) is caused by biallelic loss-of-function mutations in cystatin B (*CSTB*) (Pennacchio et al., 1996; Lalioti et al., 1997), encoding a cysteine protease (cysteine cathepsin) inhibitor with a largely unknown physiological function (Rawlings and Barrett, 1990; Turk and Bode, 1991). EPM1 is a neurodegenerative disease characterized by onset between 6 and 16 years of age, progressive drug-resistant myoclonus, epilepsy, and ataxia (Kälviäinen et al., 2008) (Fig. 1). Brain magnetic resonance imaging of EPM1 patients have revealed cortical and thalamic atrophy, as well as widespread white matter degeneration (Koskenkorva et al., 2009; Koskenkorva et al., 2012; Manninen et al., 2013). The majority of EPM1 patients are homozygous for a dodecamer repeat expansion mutation in the promoter region of *CSTB*, resulting in significantly reduced CSTB protein expression and, consequently, increased activity of cathepsins B, L and S (Rinne et al., 2002; Joensuu et al., 2007). Total loss of CSTB causes neonatal-onset encephalopathy with progressive microcephaly and hypomyelination (Mancini et al., 2016; O’Brien et al., 2017) implying an important role for CSTB in brain development (Fig. 1).

**Figure 1:**
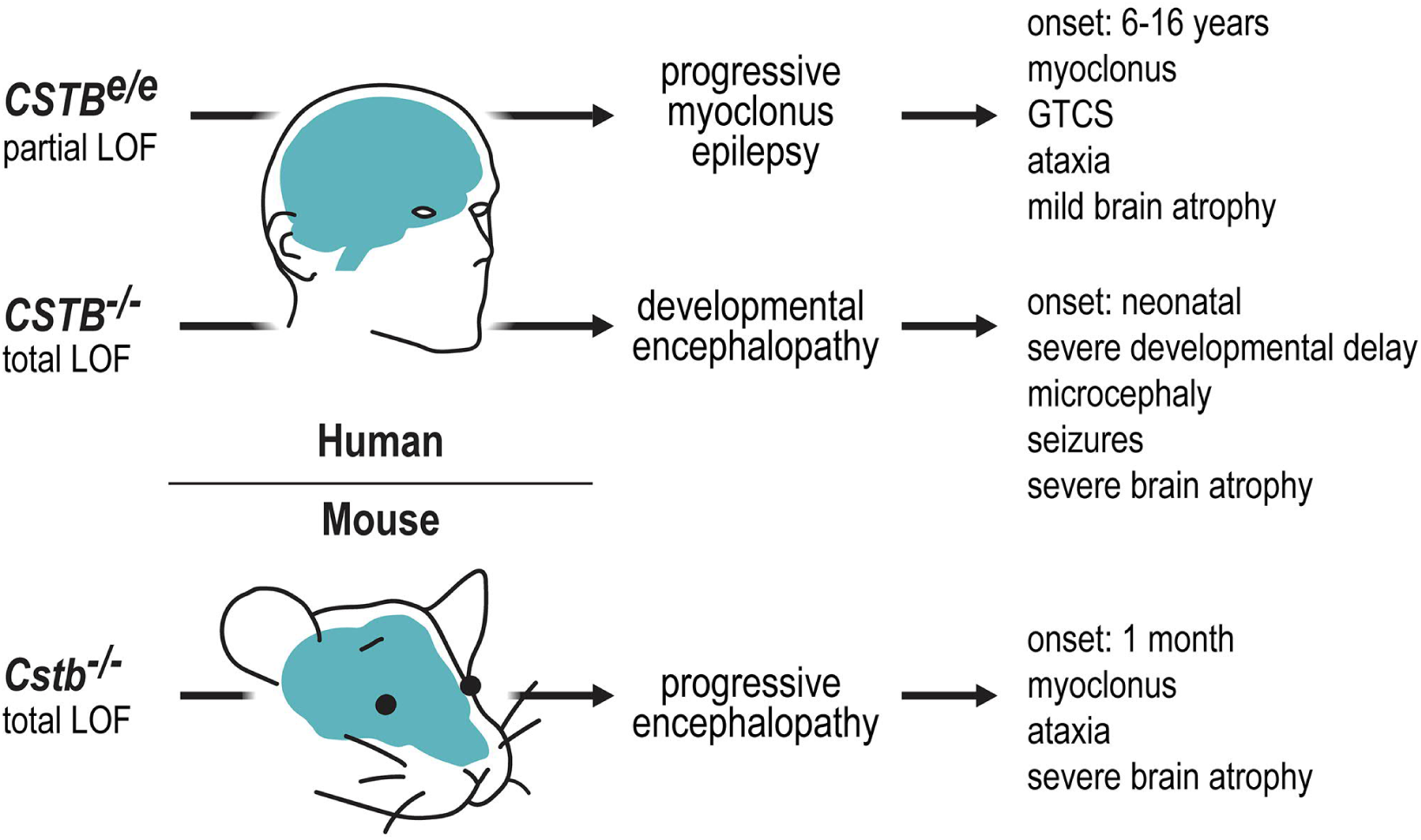
Phenotypes associated with biallelic loss of CSTB function. In human, partial loss of function (LOF) due to homozygosity for a repeat expansion in the promoter of *CSTB* gene (*CSTB*^*e/e*^) underlies progressive myoclonus epilepsy with onset in late childhood/adolescence. Total loss of function due to biallelic null mutations (*CSTB*^*−/−*^) leads to a neonatal-onset developmental encephalopathy. *Cstb* knock-out mice (*Cstb−/−*) show a neurodegenerative disorder with neuronal hyperexcitability and glial activation preceding the onset of clinical symptoms by 1 month of age. GTCS: generalized tonic-clonic seizures.

A *Cstb*-knockout (*Cstb*^*−/−*^) mouse model recapitulates key features of EPM1 including myoclonus, ataxia and progressive brain degeneration (Pennacchio et al., 1998; Shannon et al., 2002; Buzzi et al., 2012; Tegelberg et al., 2012; Manninen et al., 2013; Manninen et al., 2014) (Fig. 1). We previously demonstrated that early microglial activation and dysfunction with consequent inflammation in the *Cstb*^−/−^mouse brain precedes the onset of myoclonus and neurodegeneration (Tegelberg et al., 2012; Okuneva et al., 2015; Okuneva et al., 2016). We further showed that in the developing cerebellum, preceding microglial activation, Purkinje cells present imbalance between excitatory and inhibitory postsynaptic currents, impairment of ligand binding to GABA_A_ receptors and increased expression of GABA receptor subunit α6 and δ genes (Joensuu et al., 2014). These early alterations in GABAergic signaling, observed at post-natal day seven, suggest that CSTB-deficiency interferes with the development and/or maturation of neuronal networks.

Within the cell, CSTB localizes to the nucleus and the cytosol, where it is associated with a subset of lysosomes (Riccio et al., 2001; Brännvall et al., 2003; Alakurtti et al., 2005). In the nucleus, CSTB interacts with cathepsin L (Ceru et al., 2010; Chauhan et al., 2016), a protease with several reported nuclear functions including regulation of cell cycle progression through proteolytic processing of CDP/Cux transcription factor, stabilization of epigenetic heterochromatin markers, sperm histone degradation, and histone H3 tail clipping (Goulet et al., 2004; Bulynko et al., 2006; Duncan et al., 2008; Adams-Cioaba et al., 2011; Morin et al., 2012). Cathepsin L is one of several intracellular proteases that have emerged as epigenetic regulators through their ability to remove histone tails, particularly during cell-state transitions (e.g. Duncan et al., 2008; Khalkhali-Ellis et al., 2014; Kim et al., 2016; Yi and Kim, 2018). Histone tail clipping implies a quick and irreversible erasure of epigenetic signatures from the nucleosome, and impacts nuclear processes from gene expression to higher-order chromatin structure (Santos-Rosa et al., 2009; Nurse et al., 2013). The upstream regulation and downstream effects of this mechanism are poorly defined, which makes it one of the least understood modalities of histone post-translational modification (reviewed by Yi and Kim (2018)).

Given the function of CSTB as an endogenous inhibitor of cathepsin L, its localization in the nucleus and its predicted role in brain development, implied by both human and mouse data, we explored the hypothesis that CSTB is involved in chromatin remodeling during brain development. We found that CSTB functions as an endogenous modulator of histone H3 tail clipping through the inhibition of cysteine cathepsins B and L, and that absence of CSTB results in ectopic H3 clipping.

## RESULTS

### CSTB regulates histone H3 tail clipping during neural stem cell renewal and differentiation

Proteolytic removal of the N-terminal tail of histone H3 (histone clipping) has been observed in a number of cell types and physiologic conditions (e.g. Duncan et al., 2008; Santos-Rosa et al., 2009; Duarte et al., 2014; Khalkhali-Ellis et al., 2014; Kim et al., 2016; Melo et al., 2017; Shen et al., 2017). However, it is not known whether this mechanism is involved in neural stem cell differentiation. To address this question, we generated an *in vitro* model of neural stem cell renewal and differentiation from embryonic mouse brain (Fig. 2A). We acid-extracted histones from wild-type (*wt*) mouse neural progenitor cells (NPCs) during self-renewal (undifferentiated; uD) and differentiation (days 1, 5 and 12 post-differentiation; pD), and analyzed them by Western blotting using different antibodies against histone H3. In addition to the expected H3 band of approximately 16 kDa, we detected two faster-migrating H3 species, both of which appeared specifically upon induction of differentiation (Fig. 2B-C). An antibody against H3K4me3 only detected the intact form of histone H3, implying that the faster-migrating H3 species are the result of N-terminal truncation. The lighter of the cleaved H3 species was identified both with an antibody against the C-terminus of histone H3 (Fig. 2B-C) and with one against H3K27me3 (Fig. 2B; *Supplementary Appendix*, Fig. 2-1). These observations indicate that this cleaved H3 product has undergone N-terminal proteolysis proximal to the lysine at position 27. The heavier cleavage product was found to correspond to histone H3 truncated between alanine 21 and threonine 22 (hereinafter referred to as H3T22cl), based on specific staining with a monoclonal antibody that recognizes H3 with threonine 22 at its N-terminal end (Fig. 2B-C; *Supplementary Appendix*, Fig. 2-1).

**Figure 2:**
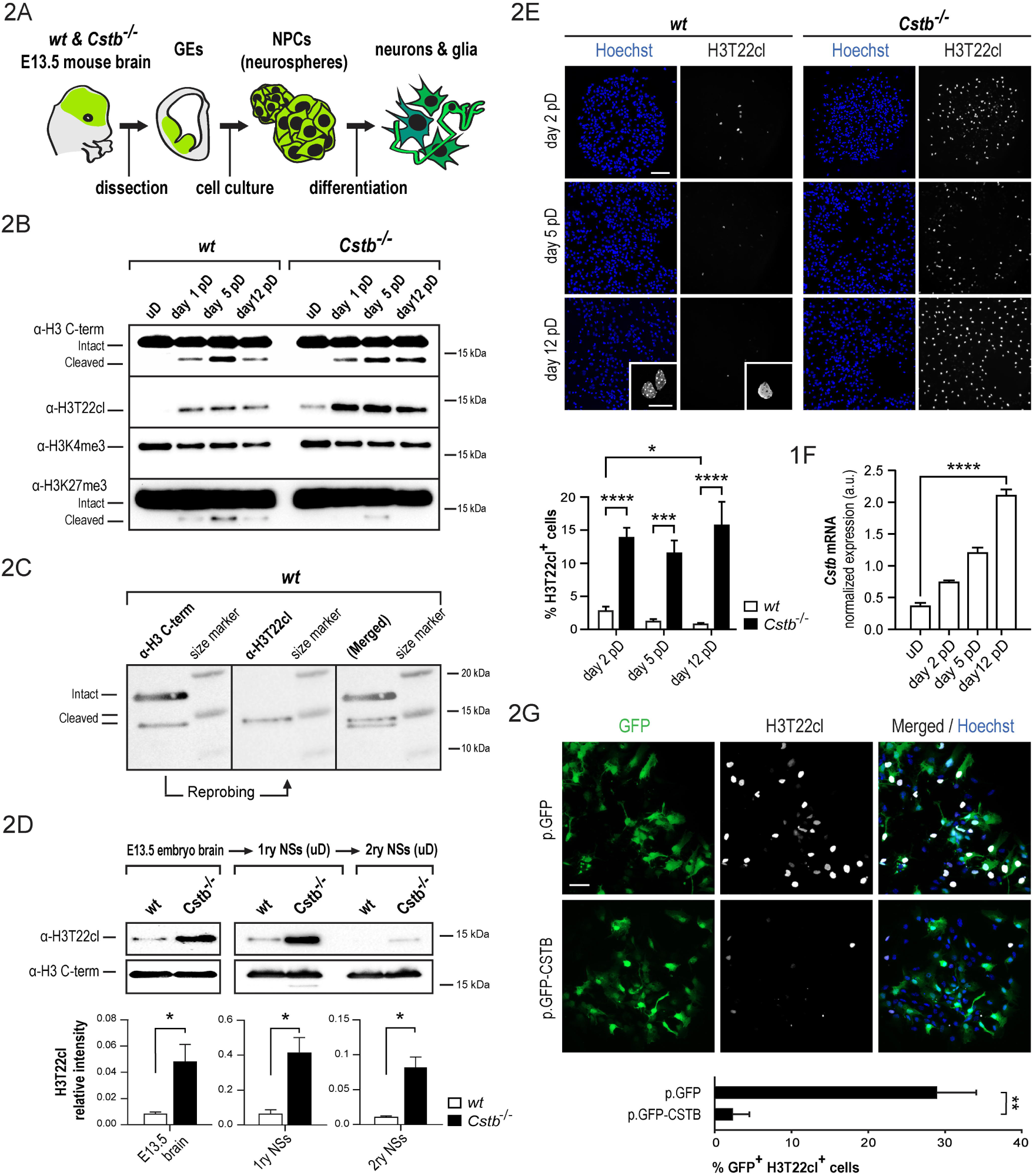
CSTB prevents ectopic clipping of the histone H3 tail in NPCs. **A:** Schematic representation of the experimental model used in this study. Briefly, NPCs derived from the ganglionic eminences (GEs) of *wt* and *Cstb*^−/−^ E13.5 mouse embryos were selectively amplified *in vitro* as neurospheres, plated for differentiation and allowed to differentiate for up to 12 days. **B:** Western blot analysis of different histone H3 modifications in histone extracts collected from *wt* and *Cstb*^−/−^ NPCs before induction of differentiation (undifferentiated, uD) and at days 1, 5 and 12 after induction of differentiation (post-differentiation, pD) using antibodies against the C-terminus of H3, H3T22cl, H3K4me3 and H3K27me3. **C:** Sequential Western blot detection of truncated products of histone H3 in *wt* NPCs at day 5 pD using antibodies against the C-terminus of H3 and H3T22cl. **D:** Western blot analysis of the relative abundance of H3T22cl in histone extracts of *wt* and *Cstb*^−/−^ E13.5 mouse embryo forebrains and cultured undifferentiated (uD) primary and secondary neurospheres (1ry and 2ry NSs). H3T22cl intensity value was normalized to that of total H3 detected with an antibody against the C-terminus of H3 (n = 3-5). **E:** Representative immunofluorescence images of differentiating *wt* and *Cstb*^*−/−*^ NPCs stained for H3T22cl and DNA using Hoechst stain. Scale bar = 100 μm. Confocal microscopy images showing a H3T22cl-positive and a H3T22cl-negative cell are displayed in the insets of *wt* day 12 pD figures. Scale bar = 20 μm. The chart depicts the proportion of H3T22cl-positive cells over total cell count. (n = 13-15). **F:** RT-qPCR analysis of *Cstb* mRNA in self-renewing and differentiating *wt* NPCs. The chart depicts *Cstb* expression normalized to mRNA level of *Rpl19* (n = 5). The results were posteriorly validated by normalization to mRNA levels of *Atp5c1* and *Ywhaz* (n = 5; data not shown). **G:** Representative immunofluorescence images of differentiating *Cstb*^−/−^ NPCs overexpressing GFP (p.GFP) or GFP-CSTB (p.GFP-CSTB). Cells were stained for H3T22cl and DNA using Hoechst stain. Scale bar = 20 μm. The chart depicts H3T22cl / GFP positive cells over GFP positive cells (n = 3). * p < 0.05, ** p < 0.01, *** p < 0.001, **** p < 0.0001.

We next investigated the potential involvement of CSTB in the regulation of histone clipping and carried out Western blot analysis of histone extracts derived from *Cstb*^−/−^ mouse NPCs during self-renewal and differentiation. We confirmed the presence of the two histone H3 cleavage products in differentiating cells (Fig. 2B). Notably, we detected the H3T22cl band also in undifferentiated NPCs (Fig. 2B), suggesting that CSTB-deficient cells undergo ectopic H3T22 clipping during self-renewal.

In order to find out whether histone H3 tail clipping also occurs *in vivo* and is not a phenomenon induced by cell culture conditions, we analyzed its presence in histone extracts from brains of *wt* and *Cstb*^−/−^ embryonic day (E) 13.5 mouse embryos by Western blotting. We detected H3T22cl in both *wt* and *Cstb*^−/−^ brain samples (Fig. 2D) with the abundance of H3T22cl being approximately five-fold higher in *Cstb*^−/−^ samples (t_(8)_ = 2.968, p = 0.017, t-test). H3T22cl in both genotypes was still readily observable in primary neurospheres after six days in culture but markedly decreased in abundance following passage to secondary neurospheres (Fig. 2D). *Cstb*^−/−^ cells showed significantly higher levels of H3T22cl at both cellular stages (primary neurospheres: t_(4)_ = 3.865, p = 0.018, t-test; secondary neurospheres: t_(6)_ = 3.510, p = 0.013, t-test; Fig. 2D), reflecting dysregulated histone clipping in the absence of CSTB taking place both *in vivo* and at successive stages of *in vitro* NPC amplification. We did not observe the lighter H3 cleavage product in embryonic tissue samples (Fig. 2D). Moreover, this cleavage product was absent in undifferentiated *Cstb*^−/−^ NPCs (Fig. 2B), implying that its regulation, contrary to that of H3T22cl, is independent from CSTB.

We focused our further analyses on characterizing the occurrence of H3T22cl on cellular level. We stained *wt* and *Cstb*^−/−^ NPCs for H3T22cl and nuclear DNA following induction of differentiation. We observed that the truncated product of histone H3 localized exclusively within the cell nucleus (Fig. 2E). We found that only a small fraction of *wt* cells was immunopositive for H3T22cl, with the number of H3T22cl^**+**^ cells decreasing along differentiation (2.8%, 1.2%, and 0.7% at days 2, 5, and 12 pD, respectively; F_(2,40)_ = 52.64, p = 0.0117, ANOVA; Fig. 2E). These data imply that H3T22 clipping in *wt* cells occurs transiently upon cellular induction. In line with these findings and the suggested role of CSTB in regulation of H3T22 clipping, RT-qPCR of *Cstb* expression showed that both self-renewing and differentiating *wt* NPCs express *Cstb* and that the mRNA level of *Cstb* significantly increases through differentiation (F_(4,16)_ = 182.9, p < 0.0001, ANOVA; Fig. 2F). Concordantly, in differentiating *Cstb*^−/−^ NPCs, the number of H3T22cl^**+**^ cells was significantly higher (5-fold or greater) than in *wt* cells (F_(5,78)_ = 15.31, p < 0.0001 at day 2 pD, p = 0.0003 at day 5 pD, p < 0.0001 at day 12 pD, ANOVA) and presented no observable decrease over time (Fig. 2E).

Finally, we sought to carry out rescue of ectopic histone clipping in CSTB-deficient NPCs by overexpressing *Cstb* transiently in *Cstb*^−/−^ NPCs with green fluorescent protein (GFP) as a transfection control. Twenty-four hours post-transfection, we triggered NPC differentiation followed by immunofluorescence staining for H3T22cl at day 1 pD. Quantification of the proportion of cells double positive for GFP and H3T22cl revealed that overexpression of CSTB is sufficient to almost completely block differentiation-associated histone clipping (t_(4)_ = 4.683, p = 0.009, t-test; Fig. 2G). Taken together, our data suggest that NPCs undergo regulated cleavage of histone H3 at position T22 during differentiation, and suggest that CSTB is inhibiting this chromatin remodeling event during self-renewal of NPCs and limiting its extent and timing during their differentiation.

### Cathepsins B and L mediate ectopic histone H3 tail clipping in CSTB-deficient neural progenitor cells

CSTB is an endogenous inhibitor of cathepsin L, a previously established H3T22 protease in mouse embryonic stem cells (Duncan et al., 2008). To determine whether cathepsin L is responsible for H3T22 clipping also in the neural cell lineage, we first cultured undifferentiated *Cstb*^−/−^ NPCs in the presence of the highly selective cysteine protease inhibitor E-64 or the cathepsin B and L inhibitor Z-FF-FMK. We then evaluated the capability of these compounds to prevent ectopic histone H3 clipping using Western blot analysis of acid-extracted histones. In *Cstb*^−/−^ cells cultured with either compound, the H3T22cl band was completely abolished (E-64: F_(2,15)_ = 174.6, p < 0.0001, ANOVA; Z-FF-FMK: F_(2,6)_ = 21.3, p = 0.003, ANOVA; Fig. 3A), suggesting that CSTB modulates H3 tail clipping through the inhibition of cathepsin L and/or cathepsin B. We next measured the enzymatic activity of cathepsins L and B in whole cell lysates of *wt* and *Cstb*^−/−^ NPCs. We detected an increase in the activity of both cathepsins in *Cstb*^−/−^ cells, implying decreased inhibition of activity in the absence of CSTB (Fig. 3B). The activity of cathepsin L was approximately 0.2-fold higher in *Cstb*^−/−^ than in *wt* control NPCs (t_(9)_ = 3.48, p = 0.007, t-test), whereas we observed an approximately 7-fold higher cathepsin B activity in *Cstb*^−/−^ compared to *wt* cells (t_(8)_ = 31.95, p < 0.0001, t-test). Finally, we cultured *Cstb*^−/−^ NPCs in the presence of the cathepsin B inhibitor CA-074. This compound strongly suppressed cathepsin B activity in the NPCs (t_(6)_ = 13.02, p < 0.0001, t-test; Fig. 3C). However, it caused only a partial decrease in the abundance of H3T22cl (F_(2,6)_ = 18.66, p = 0.018, ANOVA; Fig. 3A). Taken together, these data suggest that both cathepsins B and L are responsible for ectopic histone H3 tail clipping in CSTB-deficient NPCs.

**Figure 3:**
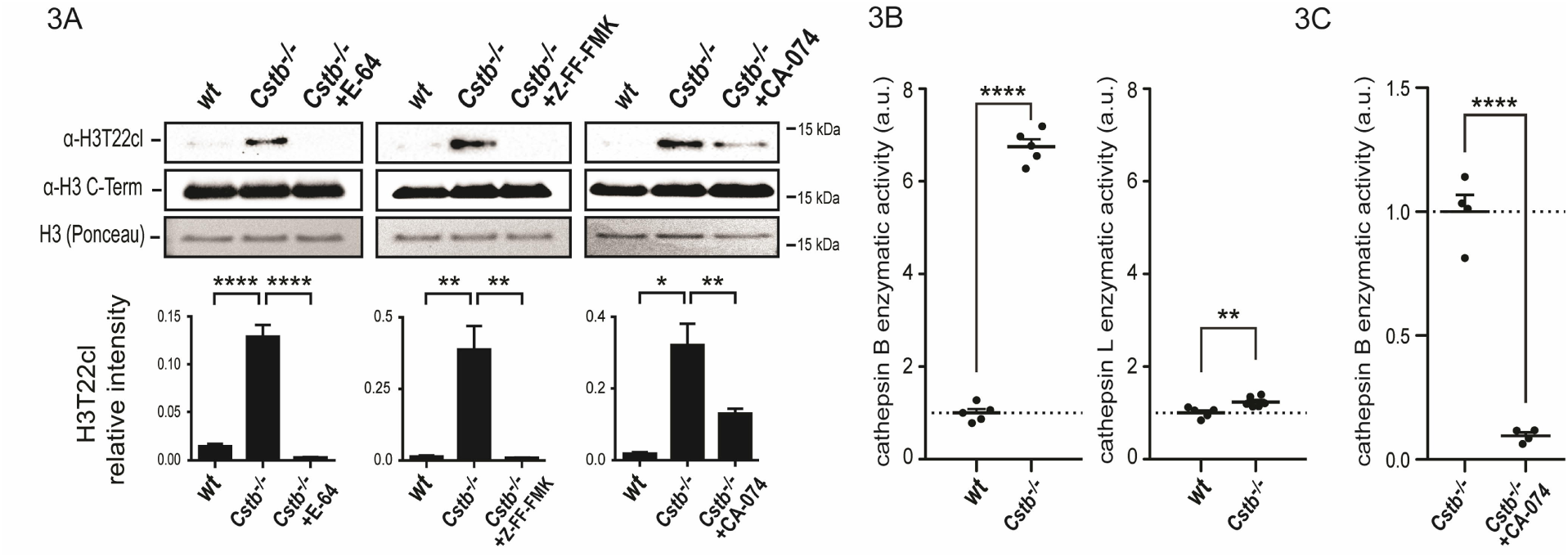
Cathepsins B and L are responsible for ectopic histone H3 clipping in *Cstb*^−/−^ NPCs. **A:** Western blot analysis and quantification of H3T22cl abundance in undifferentiated *Cstb*^−/−^ NPCs treated with E64 (n = 4-8), Z-FF-FMK (n = 3) and CA-074 (n = 3). The capacity of each compound to block H3T22 clipping was evaluated through comparison with corresponding vehicle-treated *wt* and *Cstb*^−/−^ NPCs. H3T22cl intensity value was normalized to that of total H3, detected with an antibody against H3 C-terminus. Total H3 detected with Ponceau staining is also shown. **B:** Scatter plots depicting enzymatic activity of cathepsins B and L in whole cell lysates of undifferentiated *Cstb*^−/−^ NPCs relative to *wt* NPCs levels (dotted lines) (n = 5-6). **C:** Scatter plot depicting enzymatic activity of cathepsin B in whole cell lysates of undifferentiated *Cstb*^−/−^ NPCs treated with CA-074 relative to vehicle-7treated *Cstb*^−/−^control NPCs (dotted line) (n = 4). * p < 0.05, ** p < 0.01, **** p < 0.0001.

### H3T22 clipping is most frequent in doublecortin-positive neurons

We observed that only a small fraction of differentiating *wt* NPCs are immunopositive for H3T22cl (Fig. 2D). To find out whether histone clipping is associated with cell proliferation, we first stained cells for Ki-67, the expression of which is confined to actively proliferating cells (Gerdes et al., 1983). We observed that a small proportion of H3T22cl positive cells expressed Ki-67 at day 12 pD in *wt* and at all three assessed timepoints in *Cstb*^*−/−*^ cells (*Supplementary Appendix*, Fig. 4-1), indicating that histone clipping occurs both in dividing and in non-dividing cells with a strong preference for the latter.

**Figure 4:**
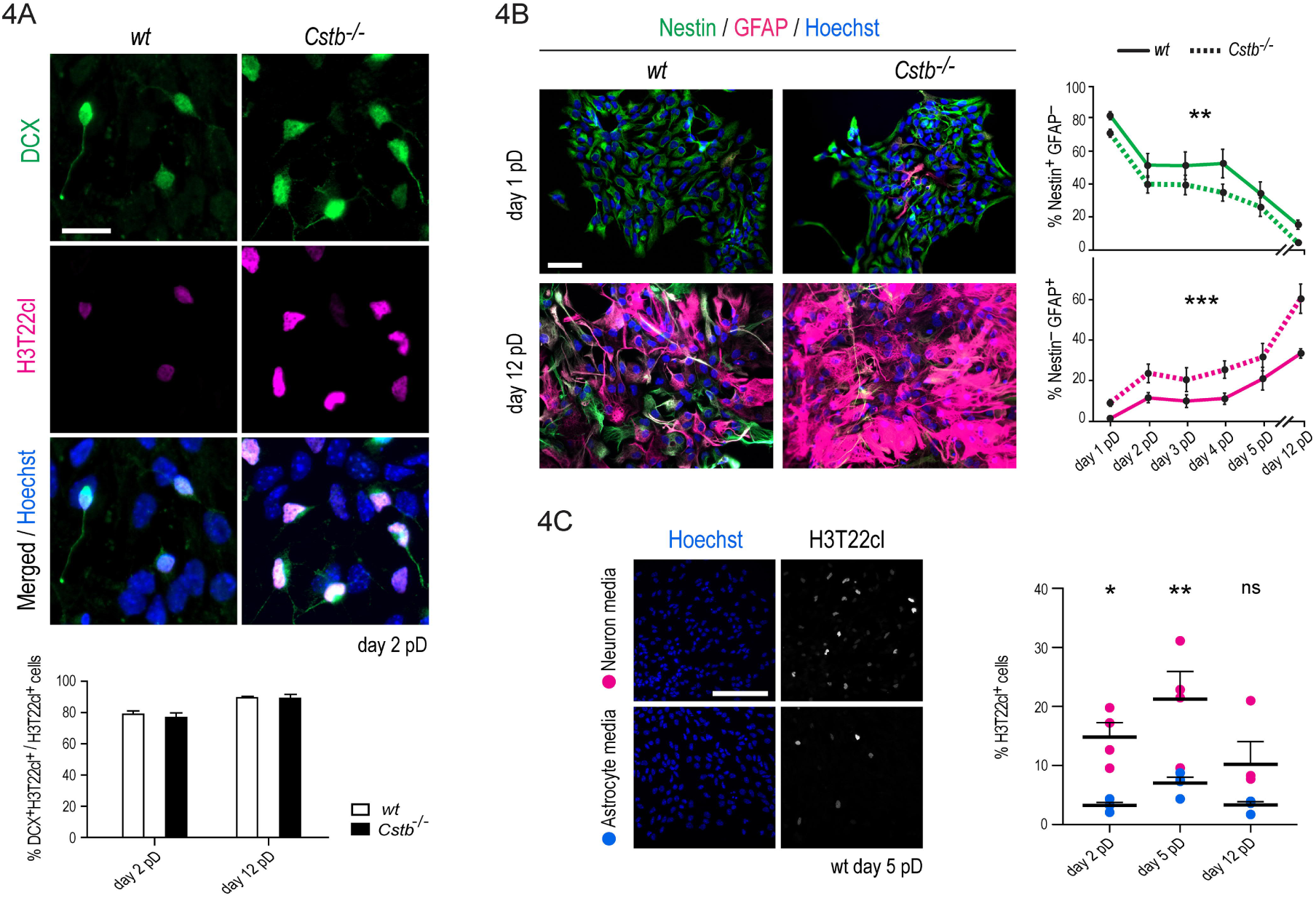
Astrogliogenesis is enhanced in differentiating *Cstb*^−/−^ NPCs yet H3T22 clipping is more prevalent in committed neurons. **A:** Representative immunofluorescence images of differentiating *wt* and *Cstb^−/−^* NPCs stained for doublecortin (DCX), H3T22cl and DNA using Hoechst stain. Scale bar = 20 μm. Bar plot depicts the proportion of DCX-positive cells among H3T22cl-positive cells (n = 6). **B:** Representative immunofluorescence image of differentiating *wt* and *Cstb*^*−/−*^ NPCs stained for nestin, GFAP and DNA. Scale bar = 50 μm. The charts depict the proportion of GFAP-negative nestin-positive cells and GFAP-positive nestin-negative cells over total cell count at six timepoints pD (n = 8). **C:** *wt* NPCs differentiated in neurogenesis-promoting media (Neuron media) or astrogliogenesis-promoting media (Astrocyte media) stained for H3T22cl and DNA using Hoechst stain. Scale bar = 50 μm. Scatter plot depicting the percentage of H3T22cl-positive cells over total cell count. (n = 4). * p < 0.05, ** p < 0.01, *** p < 0.001, ns: not significant.

To elucidate whether histone clipping occurs in all cell types or is limited to a specific cell type of the neural lineage, we next carried out immunofluorescence staining of differentiating *wt* NPCs for a number of cellular identity markers and quantified their co-expression with H3T22cl. In order to address whether CSTB deficiency causes alterations in cell type specificity of histone clipping we did the experiments parallelly in *Cstb*^−/−^ NPCs.

We first analyzed H3T22cl in differentiating neurons. Approximately 3% of both *wt* and *Cstb*^−/−^ cells at day 12 pD were positive for the neuron-specific tubulin isoform TUBB3 (TUJ1; 2.84±0.44% of *wt*, 3.14±0.56% of *Cstb*^−/−^; n = 6). Its expression was more prevalent in cells undergoing H3 tail clipping, with 10%-20% of H3T22cl^**+**^ cells being TUJ1^**+**^ neurons (*Supplementary Appendix*, Fig. 4-2) in both genotypes at all three timepoints assessed. Staining with an antibody against doublecortin (DCX) showed that approximately 9% of *wt* cells at day 2 pD, and approximately 12% at day 12 pD were committed, immature neurons with similar proportions seen in *Cstb*^−/−^ cells (*Supplementary Appendix*, Fig. 4-3). The vast majority of H3T22cl^**+**^ cells were DCX^**+**^ neurons. At day 2 pD, DCX was expressed in approximately 80% of H3T22cl^**+**^ cells and at day 12 pD in approximately 90% of H3T22cl^**+**^ cells in both genotypes (Fig. 4A).

Next, we analyzed H3T22cl in differentiating glia. During astrocyte maturation, nestin becomes progressively downregulated as cells accumulate glial fibrillary acidic protein (GFAP) (Sergent-Tanguy et al., 2006). We found that differentiating *Cstb*^−/−^ NPCs undergo transition from nestin^**+**^ GFAP^**-**^ to nestin^**-**^ GFAP^**+**^ expression at a faster rate than *wt* cells (nestin^**+**^ GFAP^−^: Chisq = 9.261, p = 0.002, linear mixed model; nestin^**-**^ GFAP^**+**^: Chisq = 16.688, p = 4.405E-05, linear mixed model), implying an impairment in the timeline of astrogliogenesis (Fig. 4B). We found that, in both genotypes, only a minority of H3T22cl^**+**^ cells were immunopositive for either nestin or GFAP, whereas the majority of H3T22cl^**+**^ cells (83% and 86% at days 2 and 12 pD, respectively) were negative for both markers (*Supplementary Appendix*, Fig. 4-4). Using immunostaining against the oligodendrocyte antigen O4, we detected late oligodendrocyte precursors in low frequency (less than 1% at days 2 and 12 pD in both genotypes). We quantified that 2%-7% of all O4^**+**^ cells were immunopositive for H3T22cl, the proportion of H3T22cl^**+**^ O4^**+**^ cells being significantly higher in CSTB-deficient conditions (40%-49%; F_(3,19)_ = 8.471, p = 0.007 at day 2 pD, p = 0.004 at day 12 pD, ANOVA; *Supplementary Appendix*, Fig. 4-5).

Taken together these data imply that all three major cell types of neural origin undergo H3T22 cleavage during differentiation and suggest that this epigenetic mechanism is most prevalent among cells committed to a neuronal fate. To gain further support to this observation, we induced differentiation of *wt* NPCs in media with altered compositions to either promote or depress neuronal differentiation. Setup efficacy was validated through quantification of GFAP^**+**^ and TUJ1^**+**^ cells at day 12 pD (*Supplementary Appendix*, Fig. 4-6). Staining for H3T22cl revealed that, compared to cells cultured in the astrocyte media, NPCs cultured in neuronal differentiation promoting media displayed a significantly higher number of H3T22cl^**+**^ cells, consistent with enhanced production of neurons (F_(5,18)_ = 7.447, p > 0.016 at day 2 pD, p = 0.003 at day 5 pD, p = 0.228 at day 12 pD, ANOVA, Fig. 4C).

### Time-resolved transcriptional changes in *Cstb*^−/−^ neural progenitor cells

It was previously shown that promoter-associated H3T22 clipping facilitates gene activation in yeast (Santos-Rosa et al., 2009). Based on these observations, we asked whether ectopic initiation of histone H3 clipping influences gene expression in *Cstb*^−/−^ NPCs. We performed RNA sequencing (RNAseq) of mRNAs isolated from *wt* and *Cstb*^−/−^ NPC populations during self-renewal and at days 1, 5 and 12 after induction of differentiation, and determined gene expression changes in the absence of CSTB.

Principal component analysis (PCA) of the transcriptomics data showed that approximately 93% of the variation between samples (PC1-PC3) originated from the process of differentiation rather than the genotype (*Supplementary Appendix*, Fig. 5-1). PC1 (77% of the variation) segregated the RNA samples chronologically, in a well-defined cluster at each timepoint assessed (Fig. 5A). The samples derived from undifferentiated NPCs formed a tight cluster, reflecting a high degree of similarity between their transcriptomes. However, PCA of this timepoint alone consistently separated *wt* and *Cstb*^−/−^ samples along the PC1 axis (32% of the variation) (Fig. 5A), indicating that the transcriptional effects of CSTB deficiency manifest already before induction of differentiation, coinciding with premature initiation of histone clipping.

**Figure 5:**
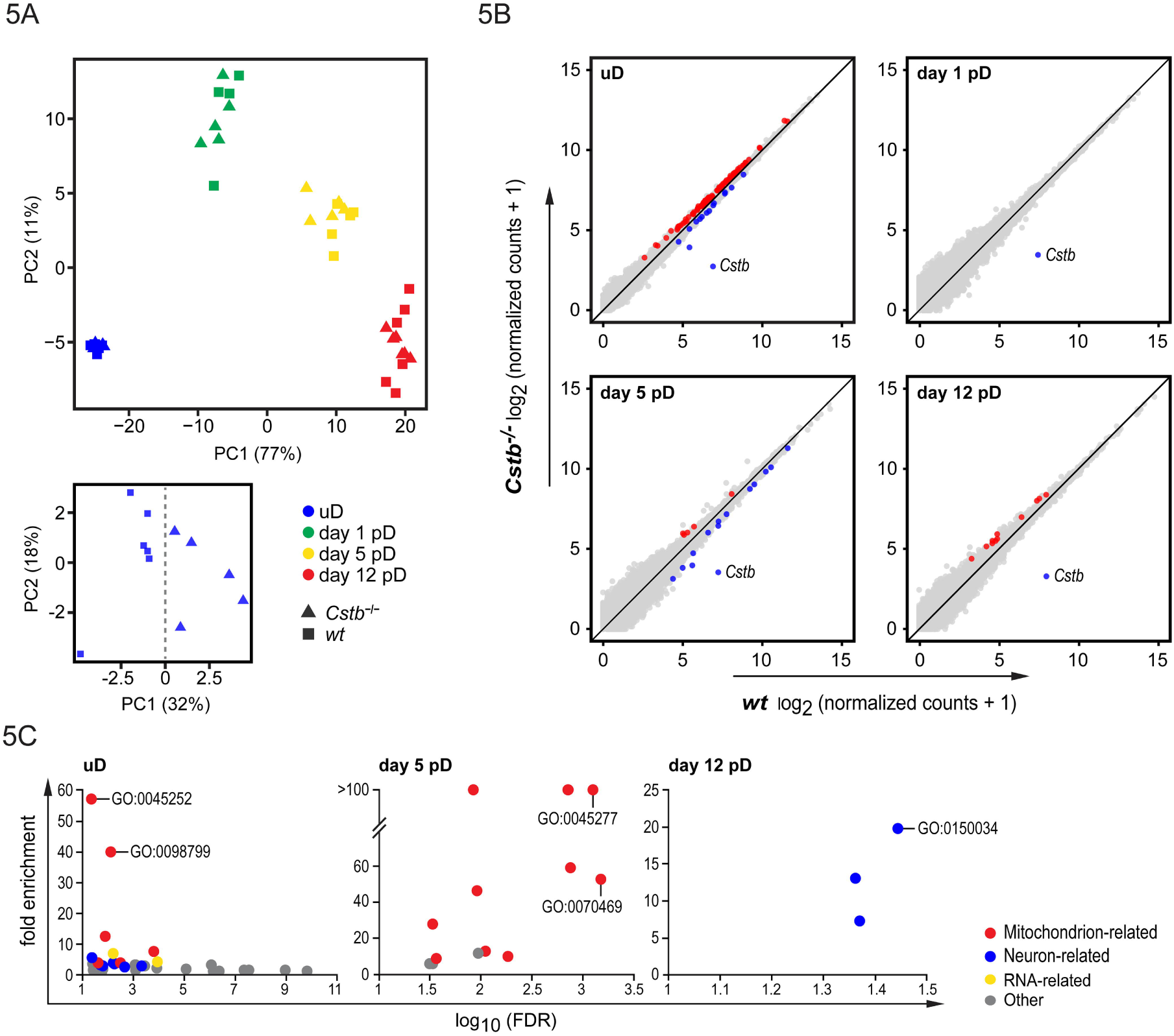
Transcriptional changes during self-renewal and differentiation in *Cstb*^−/−^ NPCs. **A:** PCA of all 42 mRNA samples analyzed by RNAseq. PCA of undifferentiated samples alone is also displayed. **B:** Scatter plots depicting all expressed genes per timepoints analyzed. X-axis and Y-axis represent mean gene expression in *Cstb*^−/−^ and *wt* mRNAs, respectively. Red and blue dots depict the significantly upregulated and significantly downregulated genes in *Cstb*^−/−^ samples, respectively. **C:** Dot charts depicting GO Cellular Component annotations with a statistically significant enrichment among differentially expressed gene datasets in *Cstb*^−/−^ at the indicated timepoints. Each category is represented as a dot plotted in relation to its significance level (base 10 logarithm of the false discovery rate (log_10_FDR), X-axis) and its fold-enrichment in the dataset (Y-axis). Colored dots represent categories related to mitochondrion, neuron and RNA metabolism as indicated. Examples of top-ranking categories: oxoglutarate dehydrogenase complex (GO:0045252), outer mitochondrial membrane protein complex (GO:0098799), respiratory chain complex IV (GO:0045277), respirasome (GO:0070469), distal axon (GO:0150034).

Comparative transcriptome analysis of undifferentiated NPCs revealed a total of 112 differentially expressed genes (DEGs) between genotypes (Fig. 5B and *Supplementary Appendix*, Table 5-1), 97 of which were upregulated in *Cstb*^−/−^ NPCs. We carried out bioinformatic analyses of these 112 genes using Gene Ontology (GO) enrichment analyses. The functionally related genes are shown in Table 5-2 (*Supplementary Appendix*). GO analyses of biological processes and molecular functions indicated enrichment of several categories related to cytoplasmic and mitochondrial protein synthesis and control of protein translation, including transcripts such as eukaryotic translation initiation factors *Eif2s3y, Eif3a, Eif4a1, Eif4B*, and *Eif5A*, and genes encoding mitochondrial ribosomal proteins, such as *Mrpl54, Mrps10*, and *Mrps15*. We also noted enrichment in RNA metabolism -related genes mostly linked to processing and splicing of pre-mRNAs, such as *Txnl4*a and *Hnrnpr*. Furthermore, GO cellular components identified 30 mitochondrion-associated genes (Fig. 5C). These included *Ndufs3, Uqcrfs1*, and *Cox6a1*, which encode proteins belonging to mitochondrial respiratory complexes I, III and IV, respectively, *Ogdh* and *Dlst*, belonging to the oxoglutarate dehydrogenase complex, and *Samm50, Tomm20*, and *Tomm40*, which encode proteins of the outer mitochondrial membrane involved in cytosolic protein import and sorting. In addition, GO terms of cellular component indicated transcripts of neuron processes and actin binding involved in regulation of cell shape and migration, such as *Pfn1* and *Dbnl*.

We did not observe other DEGs than *Cstb* at post-differentiation day 1 and next investigated the 19 and 12 DEGs in *Cstb*^−/−^ NPCs at days 5 and 12 following induction of differentiation, respectively. At day 5 pD, 14 out of the 19 DEGs were downregulated (Fig. 5B and *Supplementary Appendix*, Table 5-1). Based on GO functional classifications(*Supplementary Appendix*, Table 5-2), transcripts of electron transport chain and electron transfer activity were enriched. Accordingly, GO cellular component analysis of DEGs in *Cstb*^−/−^ mice (Fig. 5C) showed enrichment in transcripts of oxidative phosphorylation complexes I (*Ndufs2*), III (*Uqcrfs1*) and IV (*Cox6a1, Cox8a*). The 12 DEGs at day 12 pD were all upregulated in *Cstb*^−/−^ samples (Fig. 5B) and were mostly involved in neuronal maturation and integrity, as judged by enrichment in neuron-related ontologies. Half of the genes were annotated in the cellular component category “neuron projection”, and included four transcripts of the distal axon, namely *Dpysl3, Snca, Rab3a* and *Nefl*. Our dataset also contained two (*Stmn1* and *Snca*) out of three genes annotated under the biological process “regulation of thrombin-activated receptor signaling pathway”, a process involved in neuroinflammation, synaptic transmission and plasticity both in health and in disease (Ben Shimon et al., 2015; Ebrahimi et al., 2017). Notably, mutations in four of the 12 differentially expressed genes have been individually linked to human disorders of the nervous system (Polymeropoulos et al., 1997; Cuesta et al., 2002; Yang et al., 2016; Horga et al., 2017; Alsahli et al., 2018).

### Defective mitochondrial respiration in differentiating *Cstb*^−/−^ NPCs

To characterize the enrichment of functionally related gene sets in the NPC transcriptomes, we ranked all DEGs by expression fold change and carried out gene set enrichment analysis (GSEA). We found that, specifically at days 1 and 5 following induction of differentiation, genes belonging to the GO term “electron transport chain” were significantly enriched at the left pole of the ranking, indicating reduced expression in *Cstb*^*−/−*^ NPCs (Fig. 6A; q value = 0 based on 1000 permutations). To further investigate genes in this functional category, we first analyzed, using RT-qPCR in an independently generated set of samples, the expression of electron transport chain genes *Ndufs2* and *Uqcrfs1* and confirmed their downregulation in differentiating *Cstb*^*−/−*^ NPCs on day 5 pD (*Ndufs2:* t_(8)_ = 2.337, p = 0.048, t-test; *Uqcrfs1:* t_(8)_ = 2.453, p = 0.039, t-test; Fig. 6B). Next, to functionally assess electron transport activity in our cell model, we performed high-resolution respirometry of *wt* and *Cstb*^−/−^ NPCs. In undifferentiated cells, mitochondrial respiratory function was not altered between genotypes (C1&CII-linked OXPHOS: t_(3)_ = 3.131, p = 0.052, paired t-test; CIC: t_(3)_ = 0.181, p = 0.868, paired t-test; *Supplementary Appendix*, Fig. 6-1). Concordant with the GSEA results, differentiating NPCs at day 5 pD presented a significantly lower respiratory capacity in *Cstb*^*−/−*^ than in *wt* cells (C1&CII-linked OXPHOS: t_(4)_ = 5.303, p = 0.006, paired t-test; CIC: t_(4)_ = 4.276, p = 0.013, paired t-test; Fig. 6C), confirming the onset of mitochondrial dysfunction upon differentiation of CSTB-deficient NPCs.

**Figure 6:**
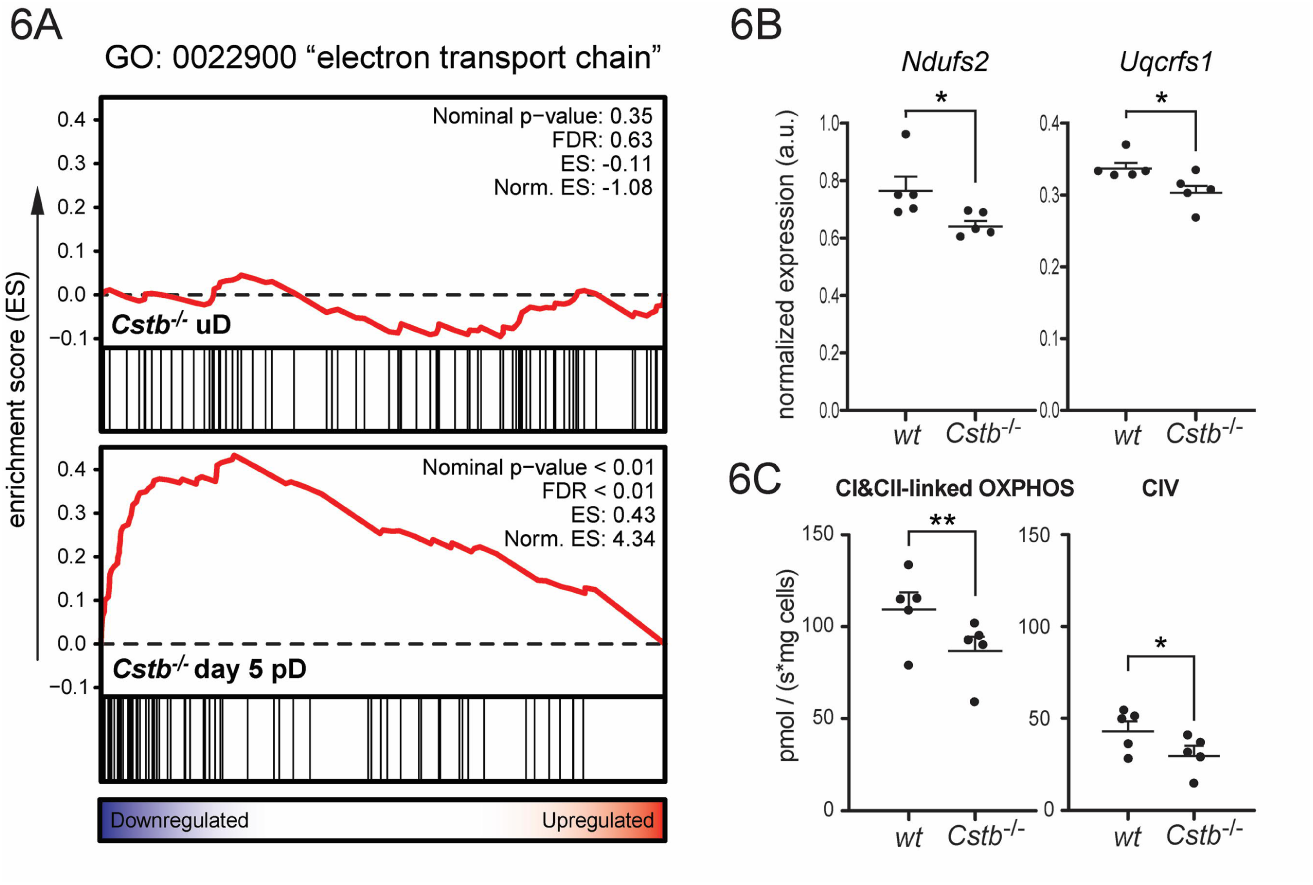
Mitochondrial dysfunction in differentiating *Cstb*^−/−^ NPCs. **A:** GSEA of differentially expressed genes in *Cstb*^−/−^ NPCs for the GO term electron transport chain. Genes are ranked from left to right from the most downregulated to the most upregulated in *Cstb*^−/−^ NPCs, respectively (one black line = one gene). **B:** RT**-**qPCR analysis of the electron transport chain genes *Ndufs2* and *Uqcrfs1* in differentiating *wt* and *Cstb*^−/−^ NPCs at day 5 pD. The chart depicts relative expression of target genes as ΔCq normalized to the mRNA levels of housekeeping genes *Atp5c1* and *Ywhaz* (n = 5). **C:** Mitochondrial respiratory capacity measurements using high-resolution respirometry in *wt* and *Cstb*^−/−^ NPCs at day 5 pD. The scatter plots depict the rate of oxygen consumption (pmol/s) by maximally coupled respiration (state III) through oxidative phosphorylation complexes I and II (CI&CII-linked OXPHOS) and individually assessed IV (CIV) normalized to protein input where each measurement represents cells derived from an individual animal (n = 5). * p < 0.05, ** p < 0.01.

## DISCUSSION

Transitions between cellular phenotypes require changes to the structure of chromatin (Dixon et al., 2015). Stem cell differentiation, for instance, involves the generation of a more restrictive chromatin state aimed at coping with the functional requirements of a specialized cellular identity (Burney et al., 2013; Zhu et al., 2013). Chromatin regulatory mechanisms are under tight control, for their synchronous action will ultimately determine the gene expression profile of the cell (Chen and Dent, 2014; Petruk et al., 2017). The proteolytic removal of histone tails, particularly of the N-terminal tail of histone H3, is one of these mechanisms. To date, clipping of the H3 tail has been observed in eukaryotic organisms from yeast to human (Santos-Rosa et al., 2009; Khalkhali-Ellis et al., 2014; Vossaert et al., 2014), has been implicated in several biological contexts such as embryonic stem cell differentiation, cellular senescence and stress response (Duncan et al., 2008; Duarte et al., 2014; Shen et al., 2017), and has been shown to impact nuclear processes from transcription to chromatin compaction (Santos-Rosa et al., 2009; Nurse et al., 2013; Duarte et al., 2014). Despite this diversity, little is known about the mechanisms upstream of histone tail removal, and its physiological significance in mammalian cells remains largely elusive (Azad et al., 2018; Yi and Kim, 2018). In the present study, we show that lack of the cysteine protease inhibitor CSTB leads to disproportionate cleavage of histone H3 in cells of the neural lineage. In humans, partial lack of CSTB is the most common underlying cause of progressive myoclonus epilepsy, EPM1 (Pennacchio et al., 1996), and complete lack of CSTB results in neonatal-onset progressive microcephaly (Mancini et al., 2016; O’Brien et al., 2017) (Fig. 1). By establishing a link between histone clipping and neurologic disease, our findings provide evidence that this epigenetic mechanism serves an irreplaceable function during brain development.

Until now, over a dozen enzymes have been shown to proteolytically process the N-terminal tail of histone H3 (Yi and Kim, 2018). However, of these only cathepsin L and an as yet unidentified serine protease involved in yeast sporulation were associated with H3 clipping at threonine 22 in living cells. We identified two seemingly independent modalities of histone H3 tail clipping in the context of neural stem cell differentiation of cultured NPCs. CSTB deficiency had no perceivable effect on the levels of histone H3 cleaved between amino acids 23 and 27, implying that this modification is not cysteine cathepsin dependent. On the contrary, CSTB deficiency impacted H3T22 proteolysis, with our data suggesting that not only cathepsin L, but also cathepsin B is responsible for this event. Considering the degree of complexity found in the mammalian nervous system, it is perhaps not unexpected to find redundancy in the machinery driving neural stem cell differentiation. Indeed, overlapping functions of these two proteases in the nervous system have previously been postulated in that genetically modified mice lacking either cathepsin L or B are viable, whereas double-deficient mice develop severe brain atrophy and die shortly after birth (Nakagawa et al., 1998; Halangk et al., 2000; Felbor et al., 2002). Interestingly, cysteine cathepsin -mediated degradation of histone H3 was recently shown to impact tissue homeostasis by contributing to chromosome segregation during mitosis (Hamalisto et al., 2020). Even if this process and the one reported by us differ significantly in their biological functions, both H3 truncation events are similarly sensitive to inhibition by CSTB, and are carried out by, at least, cathepsins B and L. The data combined support the view by Dhaenens et al. (2015) that a common group of enzymes has evolved to mediate both homeostatic and epigenetic signaling functions through histone degradation and tail clipping, respectively, in a context-dependent manner.

Antibody-based characterization of differentiating NPCs revealed that the vast majority of H3T22cl^+^ cells were immature committed neurons, which could reflect a longer temporal window for histone clipping in neurons than in glia. Lack of CSTB in our model led to enhanced astrogliogenesis, but did not seem to interfere with the rate of neuron production. In line with the latter observation, RNA sequencing of differentiating *wt* and *Cstb*^*−/−*^ NPC colonies revealed an overall similar transcriptional profile between genotypes, with only 1 (*Cstb*), 19 and 12 differentially expressed genes at days 1, 5 and 12 following induction of differentiation. Taken together, these findings indicate that *Cstb*^*−/−*^ NPCs retain their overall ability to differentiate, and suggest that the sustained cleavage of histone H3 observed in these cells does not interfere with the differentiation program. Conversely, premature initiation of histone clipping in self-renewing *Cstb*^*−/−*^ NPCs was accompanied by subtle transcriptional changes in over a hundred genes. The majority (86%) of these genes were upregulated, in accordance with previously published studies linking H3 tail clipping with transcriptional activation (Santos-Rosa et al., 2009; Kim et al., 2016). These included transcripts involved in mRNA splicing, protein translation and intracellular transport. We also detected a robust enrichment in nuclear encoded mitochondrial genes, representing approximately a quarter of the upregulated DEGs in undifferentiated cells. Interestingly, after induction of differentiation, *Cstb*^*−/−*^ cells exhibited a generalized downregulation of electron-transport-chain genes with coinciding significantly reduced mitochondrial respiratory capacity. Mitochondrial metabolism is a cornerstone of neuronal homeostasis (Oyarzabal and Marin-Valencia, 2019; Styr et al., 2019), and has profound implications for both neuronal development and disease (reviewed in Son and Han (2018)). During neural stem cell differentiation, newborn neurons undergo a metabolic transition from aerobic glycolysis to mitochondrial oxidative phosphorylation (Khacho et al., 2016). Considering our finding of histone clipping being heavily enriched among immature neurons, it is tempting to speculate that CSTB deficiency may have an impact on this neurogenesis-associated metabolic switch. Nevertheless, we demonstrate that *Cstb*^*−/−*^ NPCs manifest metabolic defects immediately upon lineage specification. Mitochondrial dysfunction is thus likely to be an important factor contributing to the early pathogenesis of CSTB-deficiency dependent neurodegeneration.

Our data are in line with the recently published findings of neurodevelopmental defects in brain organoids derived from EPM1 patients (Di Matteo et al., 2020). The authors report premature neuronal differentiation, evidenced by increased levels of DCX- and TUJ1-positive neurons, and smaller organoid sizes accompanied by decreased Ki-67 expression in neural progenitors lacking CSTB. Accordingly, we found that *Cstb*^*−/−*^ NPCs at day 12 pD express higher levels of genes involved in neuronal maturation and integrity. However, we did not detect any difference between the genotypes in the proportion of cells expressing DCX, TUJ1 or KI-67, likely reflecting the simplified nature of our cell-based model of neural stem cell differentiation. The divergent results could also signify that CSTB has a more crucial role in human than in mouse brain development. This is further supported by the observation that *CSTB* null mutations cause rapidly progressing microcephaly in humans, manifesting soon after birth and resulting in severe early functional incapacity (Mancini et al., 2016; O’Brien et al., 2017), whereas the progression of brain atrophy in *Cstb*^*−/−*^ mice is more protracted (Pennacchio et al., 1998; Shannon et al., 2002; Tegelberg et al., 2012; Joensuu et al., 2014; Manninen et al., 2014), with deterioration of motor functions appearing later during disease course.

In summary, we show that cysteine cathepsins are responsible for cleavage of the N-terminal tail of histone H3 at threonine 22 during neural stem cell differentiation. Our data suggest that the cysteine protease inhibitor CSTB plays a dual role in the regulation of this mechanism. On the one hand, it prevents the premature onset of histone clipping in self-renewing progenitors and on the other, it plays a crucial role in the termination of this chromatin-remodeling event in the late stages of differentiation. The latter finding implies that neural cells lacking CSTB are exposed to continuous removal of histone tails following terminal differentiation, a process that would directly interfere with the establishment of maturity-related epigenetic signatures in the N-terminal tail of histone H3. These results constitute the earliest phenotypic changes reported in *Cstb*^−/−^ mice, implying that epigenetic dysregulation is an initial trigger in the pathogenesis of diseases associated with CSTB deficiency.

## MATERIALS AND METHODS

### Mouse model

The *Cstb*^*−/−*^ mouse strain used in this study is 129S2/SvHsd5-*Cstb*^tm1Rm^, derived from the Jackson Laboratory strain 129-*Cstb*^*tm1Rm*^/J (stock no. 003486; https://www.jax.org/strain/003486) (Pennacchio et al., 1998). Wild type (*wt*) mice of the same age and background were used as controls. The research protocols were approved by the Animal Ethics Committee of the State Provincial Office of Southern Finland (decisions ESAVI/10765/04.10.07/2015 and ESAVI/471/2019).

### Primary cell cultures

The ganglionic eminences of E13.5 mouse embryos were mechanically dissociated and plated at 2 x 10^5^ cells/ml in 5 ml maintenance media: Dulbecco’s modified Eagle’s media (DMEM):F12 (3:1), 2% B-27, 20 ng/ml EGF (Millipore), 40 ng/ml mouse FGF-basic (PeproTech) (Ciccolini and Svendsen, 1998). For passaging, neurospheres were dissociated with TrypLE Express (Gibco) and cultured in maintenance media at 6 x 10^4^ cells/ml. Experiments were carried out with passage two neurospheres, unless otherwise specified. For differentiation, cells were plated onto poly-ornithine-coated surfaces at 3 x 10^4^ cells/cm^2^ in differentiation media; Standard: Neurobasal media (NB), 2% B-27, 2% fetal bovine serum (FBS); Neuron differentiation media, adapted from (Silva et al., 2009; Torrado et al., 2014): NB, 2% B-27, 1% FBS, 30 mM glucose, 100 ng/ml BDNF (PeproTech) and 50 ng/ml bovine serum albumin (BSA); Astrocyte differentiation media, adapted from (Brunet et al., 2004): NB, 2% B-27, 10% FBS). For protease inhibitor assays neurospheres were grown with 5 μM E-64 (in H_2_O), 10 μM Z-FF-FMK (in DMSO), 10 μM CA-074 (in DMSO) (Merck) or corresponding vehicle for four days prior to harvesting.

### Histone extraction and western blotting

Bulk histones were isolated from cells or tissue using Histone Extraction Kit (Abcam) and resolved on 4-12% Bis-Tris Plus Bolt™ gels at 4-5 ug of protein per lane in MES SDS Running Buffer (Invitrogen). Target-protein signal was detected with Horseradish Peroxidase (HRP) -conjugated secondary antibodies, followed by chemiluminescence reaction and image acquisition using ChemiDoc MP and Image Lab (v6.0; Bio-Rad). For normalization, membranes were gently stripped, re-blocked and re-probed against histone H3 C-terminus.

### Enzymatic activity assays

Substrate-based fluorometric enzymatic activity assays were performed on fresh whole cell lysates of undifferentiated NPCs using Cathepsin L Activity Assay Kit (Abcam) and Cathepsin B Activity Assay Kit (Abcam).

### Immunocytochemistry

Neurospheres were plated onto 12 mm German glass coverslips (Bellco) and cultured in differentiation media for up to 12 days. Cells were fixed with 4% PFA for 15 min at RT, followed by permeabilization with 0.1% Triton X-100 for 10 minutes. Hoechst 33342 (1:2000) was used to counterstain the nuclei.

### Antibodies

Primary antibodies used: rabbit α-H3 C-terminus (Abcam, Western blot (WB) 1:15000, RRID: AB_302613), rabbit α-H3T22cl (Cell Signaling Technologies, WB 1:1000, immunocytochemistry (ICC) 1:500, RRID: AB_2797961), rabbit α-H3K4me3 (Abcam, WB 1:2000, RRID: AB_306649), rabbit α-H3K9me3 (Abcam, WB 1:3000, RRID: AB_2797591), rabbit α-H3K27me2 (Abcam, WB 1:4000, RRID: AB_448222), mouse α-Nestin (Millipore, ICC 1:100, RRID: AB_94911), rabbit α-GFAP (Agilent Technologies, ICC 1:200, RRID: AB_10013382), rat α-GFAP (Thermo Fisher Scientific, ICC 1:500, RRID: AB_2532994), mouse α-DCX (Santa Cruz Biotechnology, ICC 1:150, RRID: AB_10610966), mouse α-βIII-tubulin (TuJ1) (Millipore, ICC 1:500, RRID: AB_2210524), mouse α-O4 (Millipore, ICC 1:200, RRID: AB_11213138), rat α-Ki-67 (Thermo Fisher Scientific, ICC 1:200, RRID: AB_10853185). For WB, HRP-conjugated swine α-rabbit (Agilent Technologies, RRID: AB_2617141) was used at 1:8000 to detect α-H3 C-terminus and at 1:5000 for all other antibody targets. For ICC, fluorescently labeled secondary antibodies (Alexa Fluor 488, 594 and 647; Thermo Fisher Scientific) were used at 1:500.

### Expression plasmids and transient transfections

To produce EGFP-CSTB fusion protein, the coding sequence of *CSTB* was cloned into an EGFP expression vector (pEGFP-N3, Clontech, GenBank: U57609). The empty EGFP vector was used as a control. C-terminal tagging of CSTB was chosen to prevent interaction with the protease recognition domain found near the N-terminus of the protein. Plasmid delivery through electroporation of passage three neurospheres was performed using Mouse neural stem cell nucleofector kit (Lonza).

## RNA sequencing

### Sample collection and library preparation

A total of 44 neurospheres from three independent cultures were harvested and lysed directly (uD samples) or plated for differentiation in round-bottom 96-well plates and lysed after 1, 5 or 12 days (pD samples). To limit the variability arising from clonal heterogeneity (Suslov et al., 2002), we focused the study on middle-sized neurospheres (approx. 200 μm, 400 cells). Cell lysis in Tris buffer (pH 7) containing 1% SDS, 10 mM EDTA and 0.2 mg/ml Proteinase K was followed by RNA extraction with RNA Clean & Concentrator kit (Zymo). 48-plex RNAseq libraries were prepared using modified STRT method with unique molecular identifiers (UMIs) (Islam et al., 2011; Islam et al., 2014). In order to remove amplification artefacts at high copy numbers, we used longer UMIs of 8 bp. To increase the coverage, the libraries were sequenced twice with Illumina NextSeq 500, High Output (75 cycles).

### STRT RNAseq data preprocessing

The raw output base call (BCL) files were demultiplexed with Picard (v2.10.10) ExtractIlluminaBarcodes and IlluminaBasecallsToSam to generate unaligned BAM files. BAM files were converted to FASTQ files with Picard SamToFastq and aligned to the mouse reference genome mm10, mouse ribosomal DNA unit (GenBank: BK000964), and ERCC spike-ins (SRM 2374) with the GENCODE (vM19) transcript annotation using HISAT2 (v2.1.0) (Kim et al., 2019). Aligned BAM files were merged with the original unaligned BAM files to generate UMI-annotated BAM files by Picard MergeBamAlignment. BAM files corresponding to each sample from different lanes and runs were merged using Picard MergeSamFiles. Potential PCR duplicates were marked with Picard Markduplicates. The resulting BAM files were processed with featureCounts (v1.6.2) (Liao et al., 2014) to assign the reads to 5’-end of genes. Uniquely mapped reads within the 5’-UTR or 500 bp upstream of protein-coding genes based on the NCBI RefSeq protein-coding genes and the first 50 bp of spike-in sequences were counted. FASTQ files after exclusion of duplicated reads have been deposited in the ArrayExpress database at EMBL-EBI under accession number E-MTAB-8934. The numbers of total and of mapped reads for each sample are summarized in Table 5-3 (*Supplementary Appendix*).

### RNAseq expression data analysis

Two samples out of the total 44 samples were excluded due to the extremely low number of mapped reads. After filtering out lowly expressed genes and spike-ins, differential expression analysis between *wt* and *Cstb*^−/−^ samples at each timepoint was performed with the ‘nbinomWaldTest’ method of an R (v3.5.2) package DESeq2 (v1.22.2) (Love et al., 2014). Genes with adjusted p-value less than 0.05 were considered as significantly differentially expressed. Principal component analysis was performed using the 500 genes with the highest variance across all the 42 samples. Gene set enrichment analysis (GSEA) was performed with GSEA (v4.0.3) using GSEAPreranked tool (Subramanian et al., 2005), where genes were preranked based on their p-values and fold changes. Mouse gene sets for Gene Ontology were downloaded from http://ge-lab.org/gskb/. GSEA plots were generated using ReplotGSEA.R with some modifications. Gene ontology terms of GO biological processes, GO cellular components and GO molecular functions significantly enriched in our datasets were calculated using Fisher’s exact test with Benjamini-Hochberg correction using web-based tools WebGestalt (version 01/14/2019) (Liao et al., 2019), Ingenuity Systems Pathway Analysis (IPA, QIAGEN, version 01-14) and the GO Ontology Resource (version 2019-12-09).

### Real-Time PCR

Total RNA was isolated with NucleoSpin RNA Plus kit (Macherey-Nagel) and reverse-transcribed with iScript cDNA synthesis kit (BioRad). RT-PCR was performed in CFX96 Real-Time System (Bio-Rad) using TaqMan Fast Advanced Master Mix (Applied Biosystems). The following TaqMan probes were used for target amplification: *Cstb* (Mm00432769_m1), *Rpl19* (Mm02601633_g1), *Uqcrfs1* (Mm00481849_m1), *Ndufs2* (Mm00467603_g1). Gene expression data was normalized to that of genes *Atp5c1* (Mm00662408_m1) and *Ywhaz* (Mm03950126_s1), chosen based on RNA sequencing data for their high and stable expression among timepoints and genotypes. Relative gene expression was calculated using the 2^−ΔCt^ method. All primers were validated and qRT-PCR assays were performed in accordance with MIQE guidelines.

### High-resolution respirometry

Oxygen consumption rates were measured using a substrate-uncoupler-inhibitor protocol on a high-resolution oxygraph (Oroboros Instruments) as previously described (Jackson et al., 2014). Briefly, neurospheres were dissociated with TrypLE express, resuspended in respiration buffer (110 mM D-sucrose, 60 mM lactobionic acid, 20 mM HEPES, 20 mM taurine, 10 mM KH_2_PO_4_, 3 mM MgCl_2_, 0.5 mM EGTA, 1g/L fatty acid-free BSA, pH7.1) and loaded at 2 x 10^6^ cells per chamber. Measurements were carried out in duplicates. Cells were digitonin-permeabilized, and oxygen consumption rates assessed in the presence of pyruvate-glutamate-malate with specific activities determined with +ADP, and +succinate (CI&CII-linked maximal coupled respiration, state III), maximal uncoupled respiration by titration of FCCP and individual assessment of complex IV by +Asc/TMPD (CIV). Specific oxygen consumption rates are expressed as pmol/second per million cells. For respirometry of day 5 pD samples, cells were washed with PBS, collected by scraping directly in respiration buffer and analyzed as described above with input normalized to amount of protein.

### Experimental design and statistical analysis

A biological replicate (n) was defined as a cell culture or tissue sample derived from an independent pool of mouse embryos (3 to 5 individuals from the same litter). In every case, at least three biological replicates were analyzed per condition (timepoint and genotype).

Immunostained cell cultures were imaged with Zeiss Axio Imager M2 microscope equipped with Axiocam 503 camera (Zeiss). Sample sizes of five or more biological replicates were used. For every biological replicate and condition, 10 random fields were captured with 20X or 40X objective. Cell numbers were quantified manually in ZEN software (Zeiss, v2.3). Total number of cells per field was defined by number of nuclei, counted manually or with automated thresholding in ImageJ software.

Data are presented as mean ± standard error of mean (represented as error bars in figure charts or displayed numerically in tables). Statistical analyses were carried out with GraphPad Prism (v5.0a) or with R Studio software. Comparisons between experimental conditions were made using Student’s t-test (unpaired unless otherwise specified) or One-way ANOVA with Bonferroni correction, depending on the experimental setup. For ICC-based characterization of astrogliogenesis, data were analyzed using a linear mixed model followed by a Wald’s chi-square test (Chisq). Differences between groups were considered statistically significant when p < 0.05.

## ACKNOWEDGEMENTS

We thank the Biomedicum Functional Genomics Unit and the Biomedicum Clinical Proteomics unit for skilled technical assistance and advice. This work has been supported by the Folkhälsan Research Foundation, the Sigrid Jusélius Foundation, the Finnish Epilepsy Association, Finska Läkaresällskapet (the Medical Society of Finland), Medicinska Understödsföreningen Liv och Hälsa r.f. (“Life and Health Medical Fund”), and Swedish Research Council. A-E.L. is a HiLIFE Fellow at the University of Helsinki. M.Y. is supported by the Scandinavia-Japan Sasakawa Foundation, the Japan Eye Bank Association, the Astellas Foundation for Research on Metabolic Disorders, and the Japan Society for the Promotion of Science Overseas Research Fellowships. S.K. is supported by Jane and Aatos Erkko Foundation and Swedish Research Council. The computations in this work were performed on resources provided by SNIC through Uppsala Multidisciplinary Center for Advanced Computational Science (UPPMAX) under Project SNIC 2017/7-317.

The authors declare no competing financial interests.

## SUPPLEMENTARY APPENDIX

**Figure 2-1:**
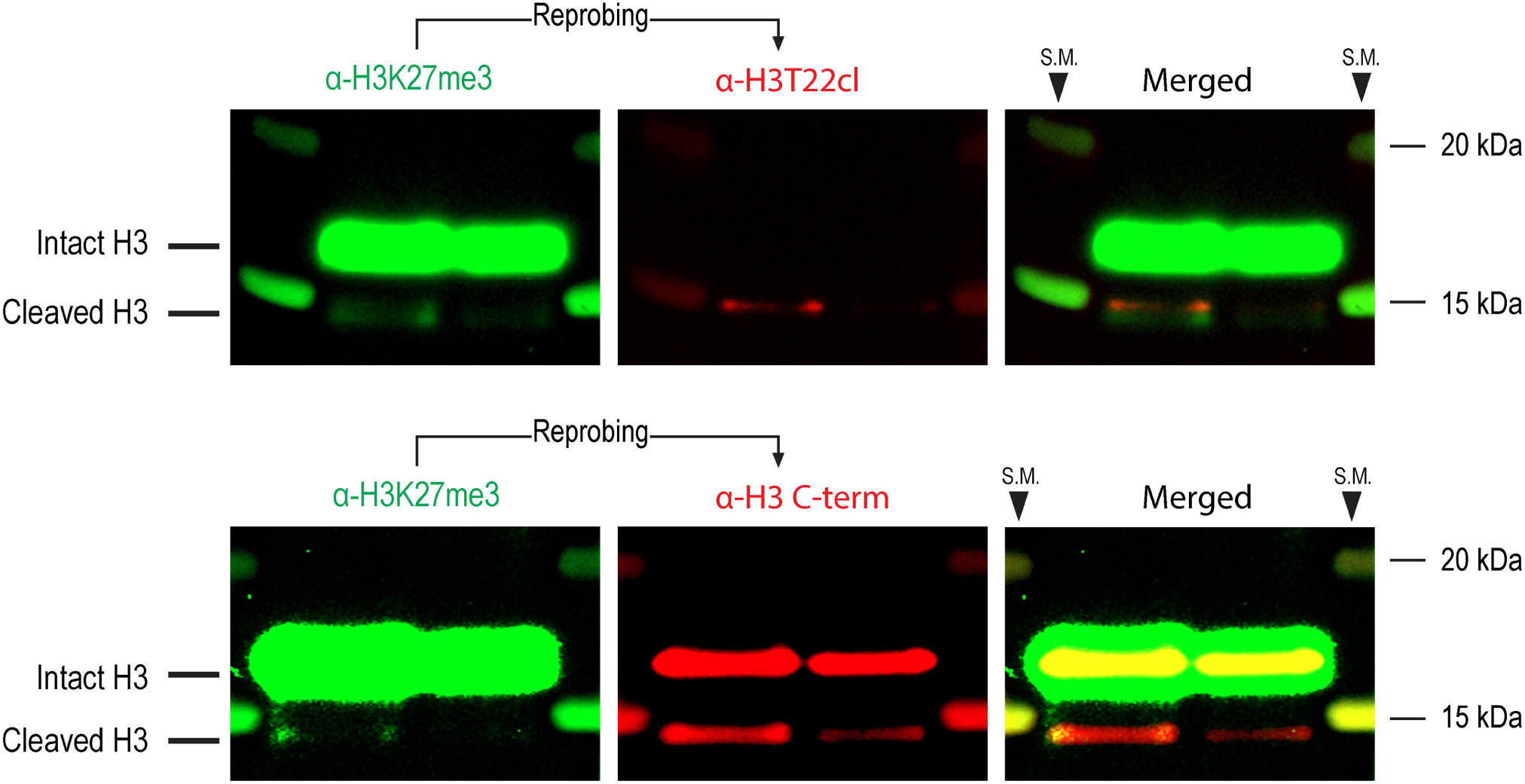
Both differentiation-associated H3 cleavage products detected in *wt* NPCs are proximal to amino acid 27. An antibody against H3K27me3 recognizes both cleavage products of histone H3 observed upon differentiation of cultured NPCs. Western blot analysis of histone extracts from differentiating *wt* NPCs at day 5 pD. Membranes were probed sequentially with antibodies against H3K27me3 and either H3T22cl (top) or H3 C-terminus (bottom). Lanes from left to right: Protein size marker / 4 μg histone / 2 ug histone / Protein size marker.

**Figure 4-1:**
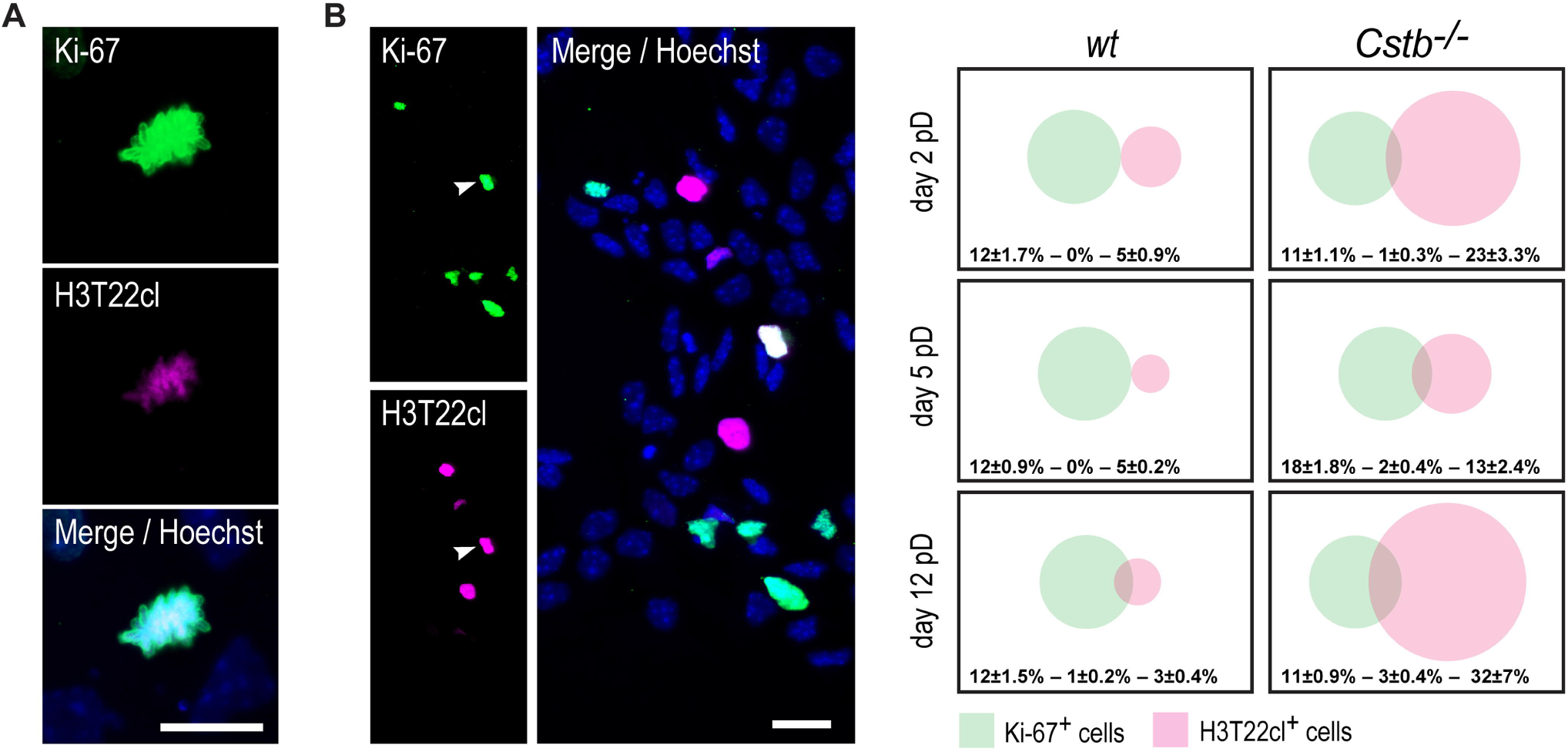
H3T22 clipping is most prevalent among non-dividing cells. Immunofluorescence images of differentiating *wt* NPCs at day 12 pD stained for Ki-67, H3T22cl and DNA (Hoechst). **A:** Ki-67 / H3T22cl positive cell at the prophase of mitosis. Scale bars = 20 μm. **B:** Representative example of the staining with one Ki-67 / H3T22cl positive cell (arrowhead). Venn diagrams depict overlap between Ki-67-positive and H3T22cl-positive cells in relation to the total cell count at the indicated timepoints, both in *wt* and in *Cstb*^−/−^ cells. Numerical values indicate average proportions ± standard error of the mean (n = 6).

**Figure 4-2:**
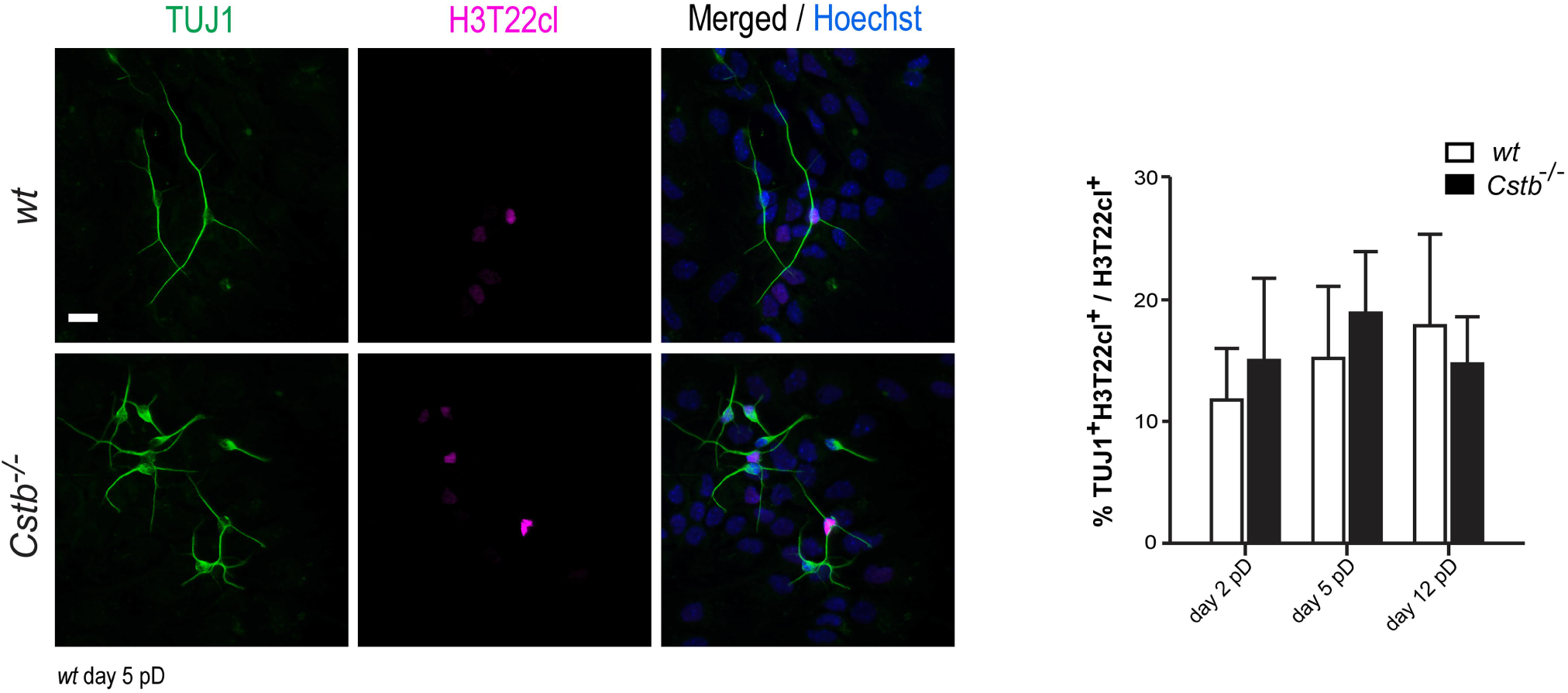
H3T22 cleavage is enriched in TUJ1-expressing neurons. Representative immunofluorescence images of differentiating *wt* and *Cstb*^−/−^ NPCs stained for TUJ1, H3T22cl and DNA (Hoechst). Scale bar = 20 μm. Bar plot depicts the proportion of TUJ1/H3T22cl -positive cells over total H3T22cl-positive cell count (n = 6).

**Figure 4-3:**
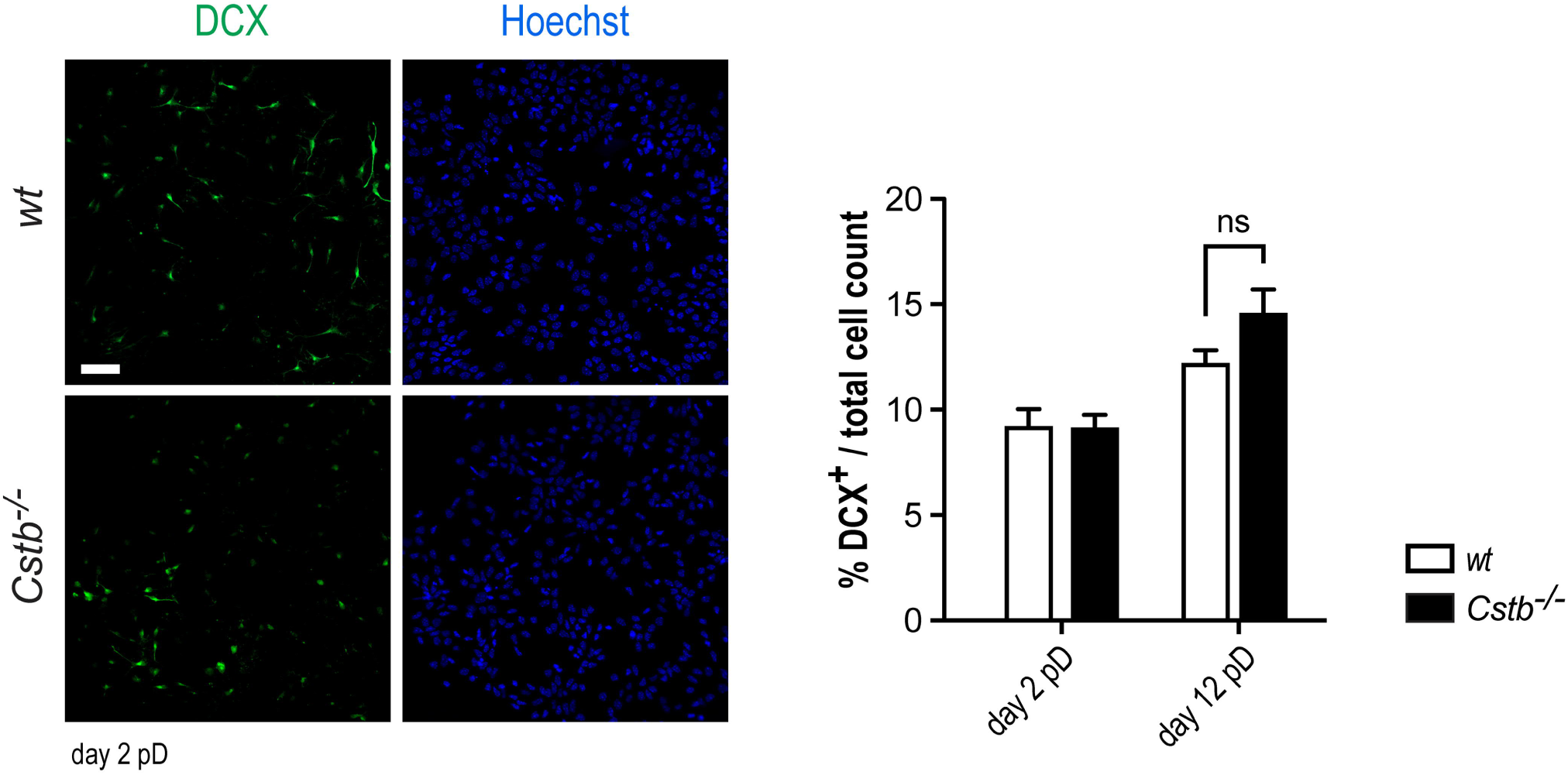
Both *wt* and *Cstb*^−/−^ NPCs give rise to DCX^+^ neurons in similar proportions. Representative immunofluorescence images of differentiating *wt* and *Cstb*^−/−^ NPCs stained for doublecortin (DCX) and DNA (Hoechst). Scale bar = 50 μm. Bar plot depicts the proportion of DCX-positive cells over total cell count (n = 5). ns: not significant.

**Figure 4-4:**
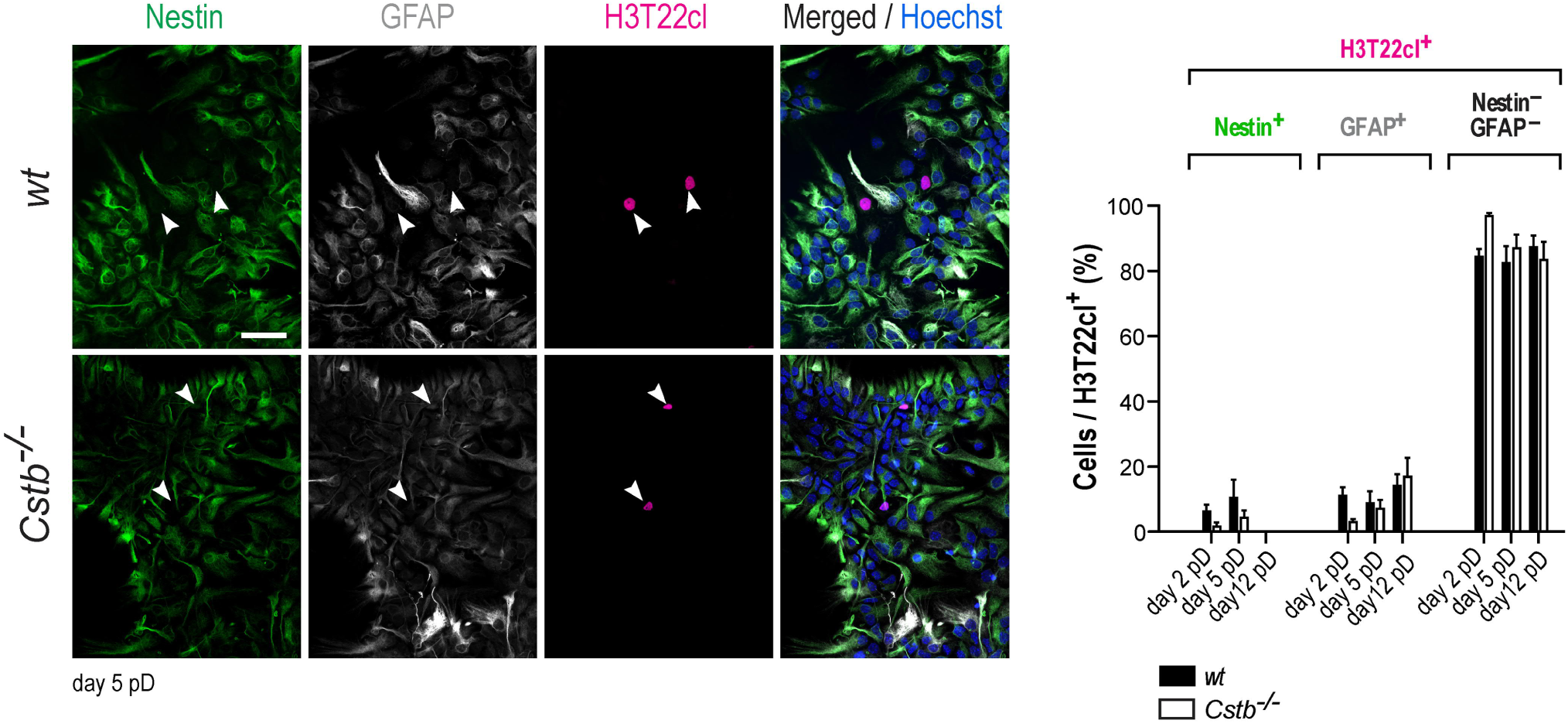
The majority of H3T22cl-positive cells do not express nestin or GFAP. Representative immunofluorescence images of differentiating *wt* and *Cstb*^−/−^ NPCs stained for nestin, glial fibrillary acidic protein (GFAP), H3T22cl and DNA (Hoechst). Arrowheads indicate the locations of H3T22cl-positive cells. Scale bar = 50 μm. The chart depicts the proportion of nestin-positive, GFAP-positive and nestin/GFAP-negative cells among total H3T22cl-positive cell population (n = 3-6).

**Figure 4-5:**
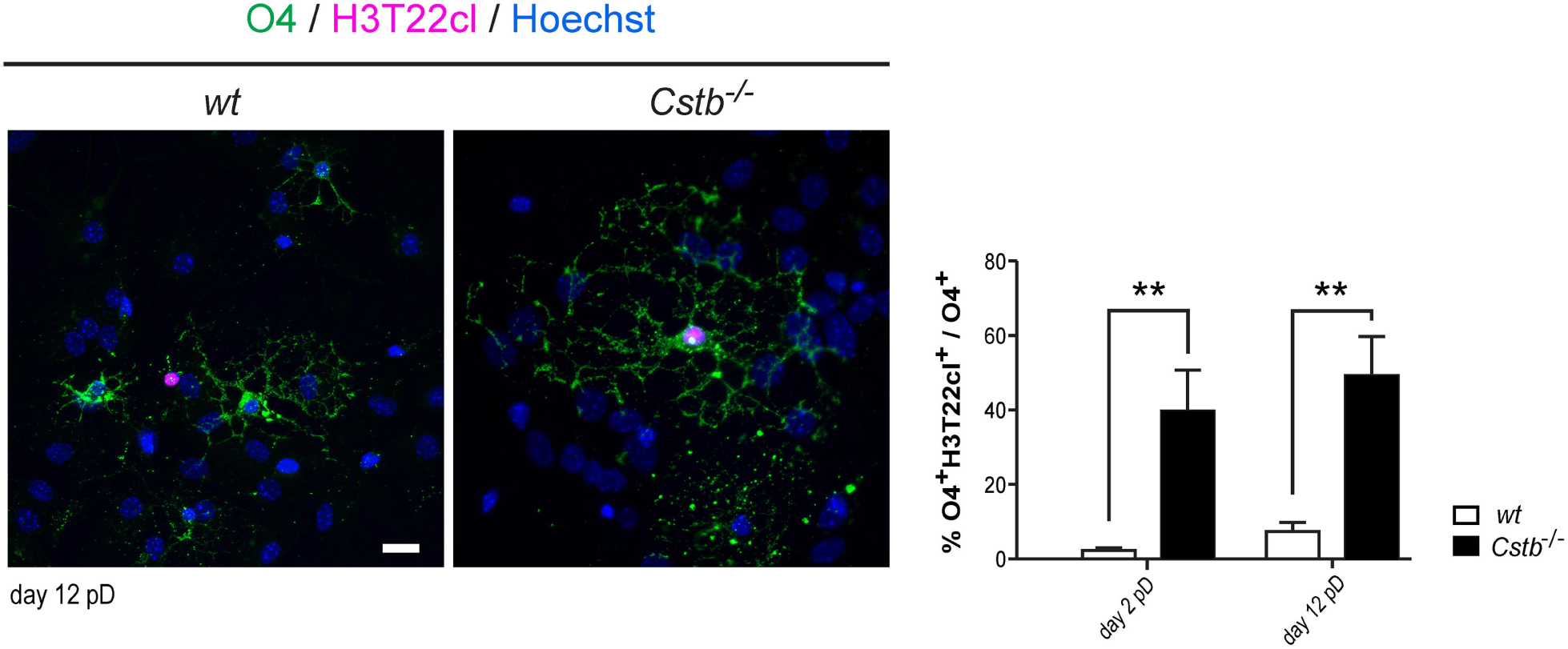
Histone H3 tail clipping takes place in cells of the oligodendrocyte lineage. Representative immunofluorescence images of differentiating *wt* NPCs stained for oligodendrocyte surface antigen O4 and H3T22cl. Scale bar = 20 μm. The chart depicts the proportion O4 / H3T22cl -positive cells over O4-positive cell count (n = 6). ** p < 0.01.

**Figure 4-6:**
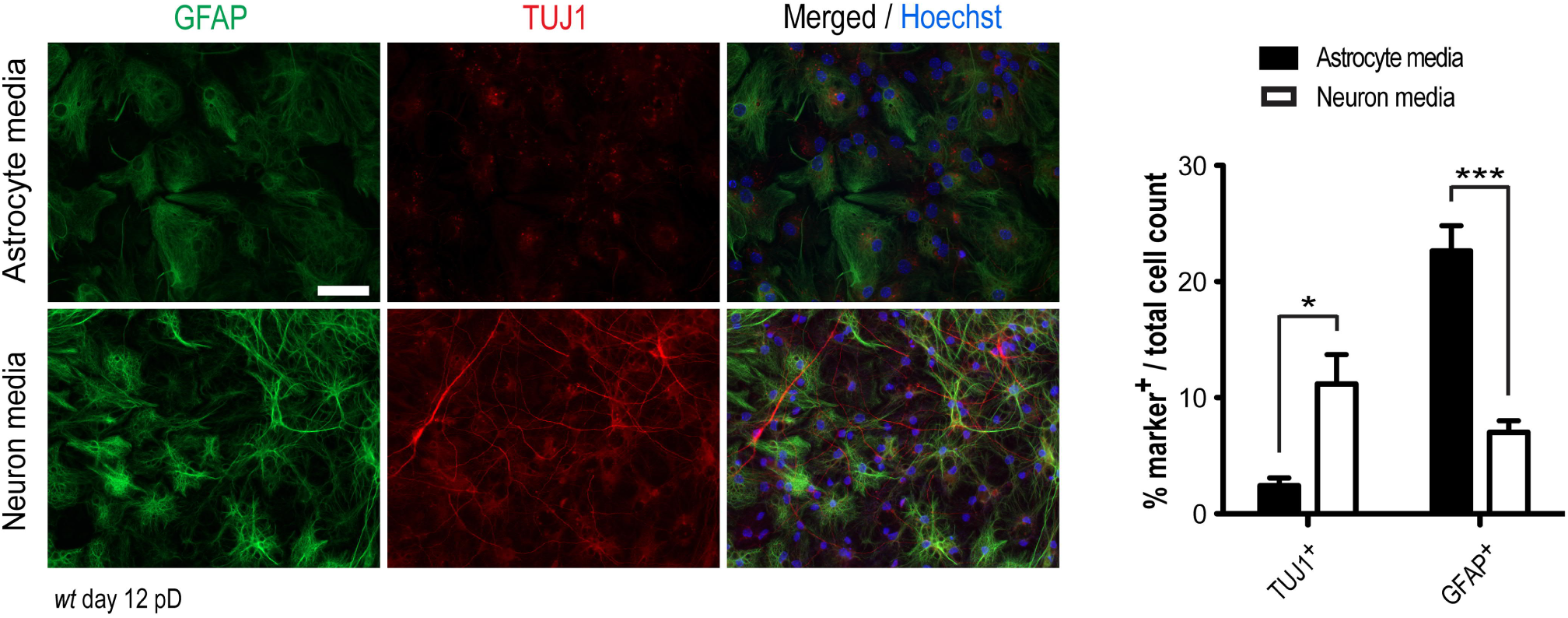
Changes in the relative production of neurons and astrocytes upon NPC differentiation in “astrocyte media” and “neuron media”. Representative immunofluorescence images of differentiating *wt* NPCs cultured for 12 days under neurogenesis-promoting conditions (neuron media) and astrogliogenesis-promoting conditions (astrocyte media) and stained for GFAP and TUJ1. Scale bar = 50 μm. Bar plot depicts the proportion of GFAP-positive and TUJ1-positive cells over total cell count at day 12 pD (n = 4-5). * p < 0.05, *** p < 0.001.

**Figure 5-1:**
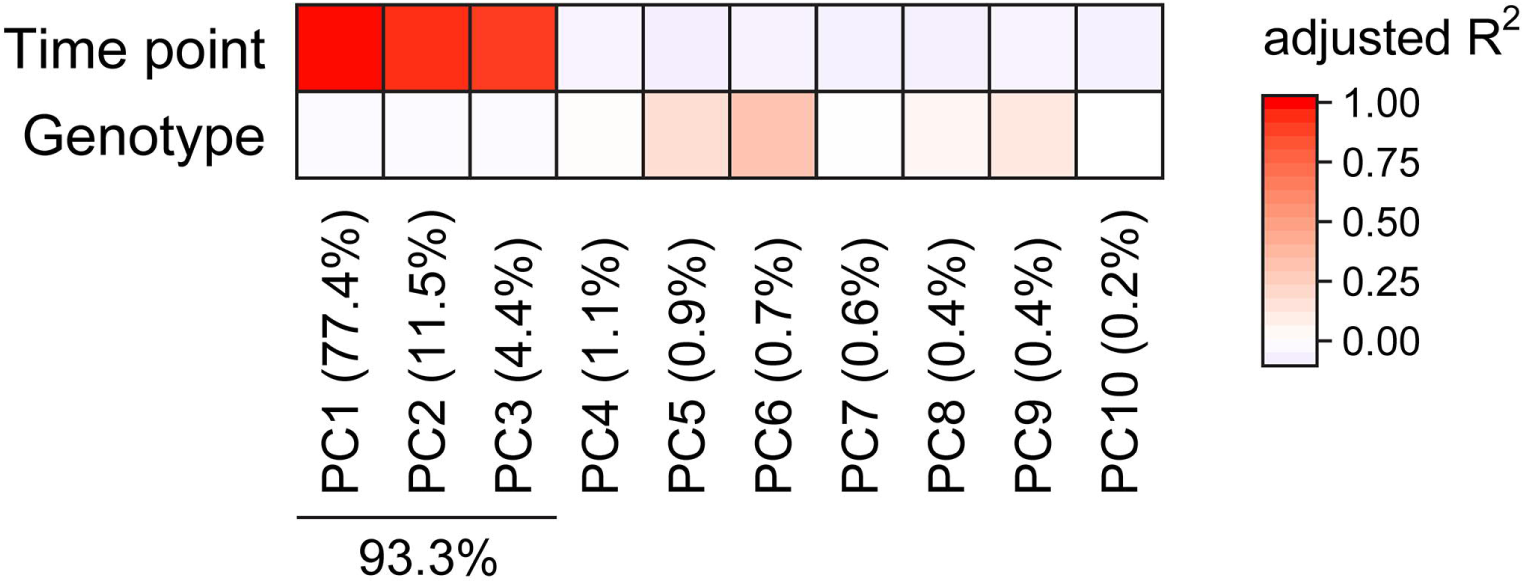
Transcriptional divergence among RNA-sequencing samples arises primarily from the process of differentiation. Heat map showing contribution of each of the first ten principal components of the RNA-sequencing data to sample variability arising from the timepoints (uD, day 1 pD, day 5 pD and day 12 pD) and genotypes (*wt* and *Cstb*^−/−^) analyzed. Colors represent adjusted R-square in linear regression. Proportion of variance explained by each principal component is also shown. Note that PC1 to PC3 account for sample heterogeneity arising exclusively from maturation status, being 93.3% of the overall sample-to-sample variation.

**Figure 6-1:**
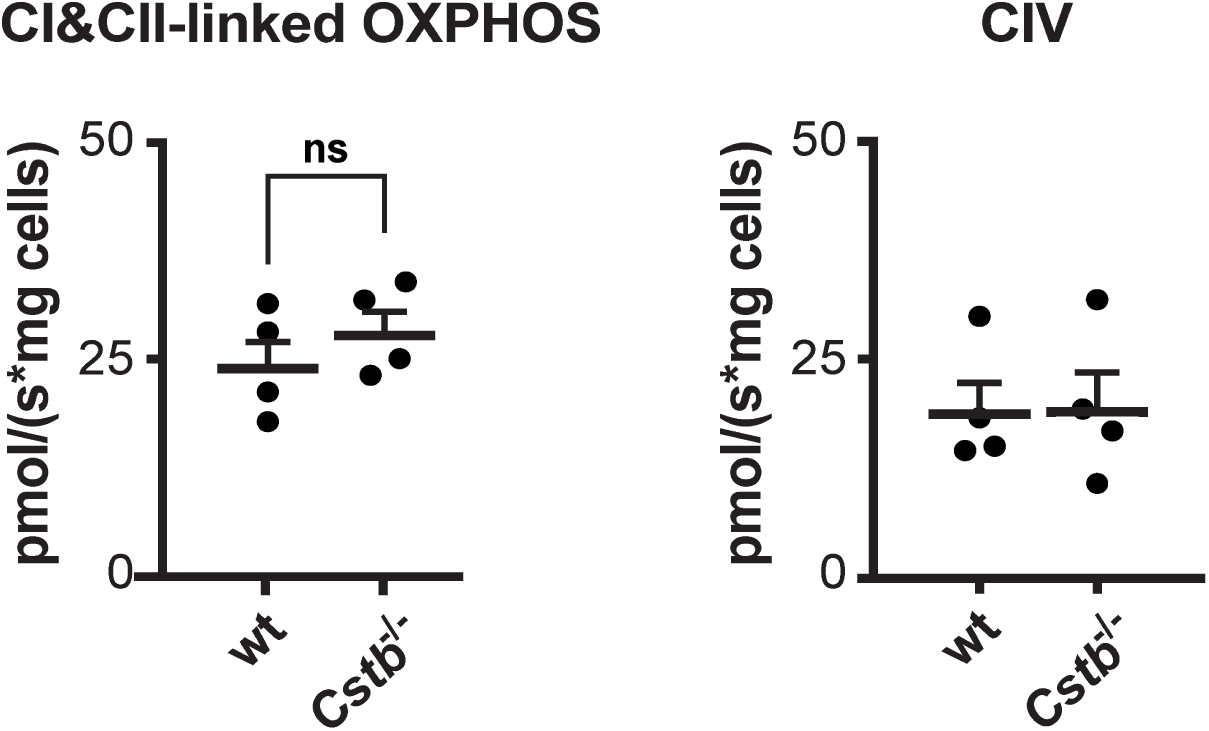
*Cstb*^*−/−*^ NPCs show statistically normal mitochondrial respiratory capacities before induction of differentiation. Mitochondrial respiratory capacity measurements using high resolution respirometry in undifferentiated *wt* and *Cstb*^−/−^ NPCs. The scatter plots depict the rate of oxygen consumption (pmol/s) by maximally coupled respiration (state III) through oxidative phosphorylation complexes I and II (CI&CII-linked OXPHOS) and individually assessed IV (CIV) normalized to input cell number (ttest; n = 4). ns: not significant.

**Table 5-1:** Differentially expressed genes (DEGs) in undifferentiated and differentiating *Cstb*^−/−^ NPCs at days 1, 5 and 12 pD.

**Table 5-2:** Gene Ontology (GO) term enrichment analysis of differentially expressed genes in *Cstb*^−/−^ NPCs for GO annotations of biological process, cellular component and molecular function. Analyses were carried out in geneontology.org (GO Ontology database version 2019-12-09).

**Table 5-3:** RNA sequencing summary.

